# Ecogenomics of key prokaryotes in the arctic ocean

**DOI:** 10.1101/2020.06.19.156794

**Authors:** Marta Royo-Llonch, Pablo Sánchez, Clara Ruiz-González, Guillem Salazar, Carlos Pedrós-Alió, Karine Labadie, Lucas Paoli, Tara Oceans Coordinators, Samuel Chaffron, Damien Eveillard, Eric Karsenti, Shinichi Sunagawa, Patrick Wincker, Lee Karp-Boss, Chris Bowler, Silvia G Acinas

## Abstract

The Arctic Ocean is a key player in the regulation of climate and at the same time is under increasing pressure as a result of climate change. Predicting the future of this ecosystem requires understanding of the responses of Arctic microorganisms to environmental change, as they are the main drivers of global biogeochemical cycles. However, little is known about the ecology and metabolic potential of active Arctic microbes. Here, we reconstructed a total of 3,550 metagenomic bins from 41 seawater metagenomes collected as part of the Tara Oceans expedition, covering five different Arctic Ocean regions as well as the sub-Arctic North Atlantic Ocean and including various depths and different seasons (spring to autumn). Of these bins, 530 could be classified as Metagenome Assembled Genomes (MAGs) and over 75% of them represented novel species. We describe their habitat range and environmental preferences, as well as their metabolic capabilities, building the most comprehensive dataset of uncultured bacterial and archaeal genomes from the Arctic Ocean to date. We found a prevalence of mixotrophs, while chemolithoautotrophs were mostly present in the mesopelagic Arctic Ocean during spring and autumn. Finally, the catalogue of Arctic MAGs was complemented with metagenomes and metatranscriptomes from the global ocean to identify the most active MAGs present exclusively in polar metagenomes. These polar MAGs, which display a range of metabolic strategies, might represent Arctic sentinels of climate change and should be considered in prospective studies of the future state of the Arctic Ocean.

## Introduction

The Arctic is under increasing pressure from climate change and growing interests in economic opportunities (e.g., natural resources such as oil and gas, tourism, etc.)^1^. Arctic microorganisms are the foundation of the marine food web, so we need to understand how they adapt and thrive, as well as to forecast their fate in a future ocean impacted by anthropogenic change. In addition, the predicted invasion of the Arctic Ocean by species from lower latitudes due to temperature increases might alter the dynamics of the entire marine ecosystem, from microbes to large animals^2^.

The Arctic Ocean’s ecosystem is subject to extreme seasonal variations (i.e., solar radiation, ice cover, temperature) and receives large inputs of freshwater rich in dissolved and organic material from rivers, as well as inflowing waters from the Pacific and the Atlantic Oceans^3^. Organisms inhabiting the upper water column thus have to adapt to a highly dynamic environment^4^. Photosynthetic primary production occurs mostly during the spring and summer seasons when light availability and increased temperatures enhance phytoplankton growth, with blooms forming under the ice cover and in the marginal ice zone^5^. Such blooms trigger a succession of bacterial populations, mostly heterotrophs from the phyla Bacteroidetes and Proteobacteria^6^. The vertical flux of organic matter during spring and summer is mainly derived from phytoplankton blooms and zooplankton fecal pellets and is highly dependent on ice-melting^7^. During winter, the lack of light makes productivity almost negligible, resulting in very low vertical carbon export from surface layers^8,9^. As photosynthesis is limited, heterotrophic bacteria and protists become the dominant players in the ecosystem^10,11^. During the polar night, other metabolisms such as mixotrophy^12,13^ and chemolithoautotrophy^14–16^ increase in importance among specific taxa of archaea and bacteria.

The Arctic Ocean can be divided into eight regions based on different features of ecological significance (AMAP/CAFF/SDWG 2013 - Identification of Arctic Marine Areas of Heightened Ecological and Cultural Significance). The record of prokaryotic diversity in such regions is scarce and generally limited to local surveys, mostly dependent on PCR amplicon sequencing and other molecular approaches involving fluorescence in situ hybridization and/or microautoradiography^17–20^. Only a few studies have attempted to assess the biodiversity of microbes across the different Arctic Ocean regions, such as the Arctic Ocean Survey (AOS) ^21^ or the International Census of Marine Microbes (ICoMM)^22^. Additionally, many countries in direct contact with Arctic waters have carried out sampling both for microbial diversity at local scales and in long term monitoring programs^23^. Previous studies have provided an overview of the microbial taxonomic diversity in the Arctic^16,20,24– 26^, including functions relevant to the ecosystem, like nitrification processes^15,27^, heterotrophy^13,21,24^ or photoheterotrophy^28,29^. Finally, the uniqueness of polar environments has been evidenced when studying global biogeographical patterns by means of amplicon sequencing^30^ or metagenomics and metatranscriptomics^31^. Recent technological advances such as the reconstruction of genomes from metagenomes are allowing to go beyond the community level and explore the functional capabilities of specific taxa. For example, metagenomic assembled genomes (MAGs) of polar origin have revealed the global biogeography of SAR11^32^ and the presence of certain genomes with the potential for carbon fixation and metabolism of nitrogen and sulfur^33^. Nevertheless, a thorough analysis of key active microbial players including their habitat and metabolic preferences in the Arctic Ocean is lacking.

The *Tara* Oceans expedition performed a holistic survey of the Arctic Ocean’s marine microbial diversity^34,35^ from May to October 2013, aiming to cover as much environmental variability within the Arctic Ocean as possible. In this study, we have built 3,550 genomic bins using the 41 prokaryote-enriched (0.22-3μm) metagenomes collected from photic to mesopelagic depths during the expedition. These bins collectively constitute a large fraction of the Arctic prokaryotic diversity detected by metagenomics and metatranscriptomics and, due to their quality scores^36^, 530 of them can be considered MAGs (Metagenome Assembled Genomes). We have performed an exhaustive pan-Arctic eco-genomic approach to the study of key uncultured Arctic prokaryotic MAGs, exploring their expression patterns, habitat preferences and metabolic potential. We identified polar sentinel genomes by selecting those MAGs found exclusively in polar metagenomes and highly transcribed within their habitat range category in Arctic samples, as a means to serve as a baseline for future monitoring of the state of the Arctic Ocean.

## Results and Discussion

### CO-ASSEMBLY AND TRENDS OF PROKARYOTIC ARCTIC BINS

The 41 metagenomes (from 0.22-3 μm size fractions) used here cover a broad range of environmental and spatio-temporal conditions in the Arctic (**Figure 1A**,**B**). We consider three different ocean layers (surface, deep chlorophyll maximum (DCM) and mesopelagic) in five Arctic Ocean regions (four stations in the Atlantic Arctic, five stations in the Kara-Laptev Sea, four stations in the Pacific Arctic, one station in the Arctic Archipelago and four stations in the Davis-Baffin) and two stations in the sub-Arctic North Atlantic. The sampling period encompasses spring, summer and autumn conditions with a wide range of temperatures (from −1.7 to 11.1 °C), sea ice conditions and photoperiods. Different assembly strategies have been used in the recovery of environmental genomes from metagenomes, like assemblies of single samples^37^ or co-assembly and binning of geographically delimited samples^38^. Here we chose to co-assemble pools of samples that were most similar in their community-level taxonomic composition (assessed with 16S miTags and NMDS with 100 iterations and stress value of 0.08), as a way to obtain a less redundant set of bins with higher genome completeness, and binned all resulting contigs in one batch (**Figure S1**).

**Figure 1.**
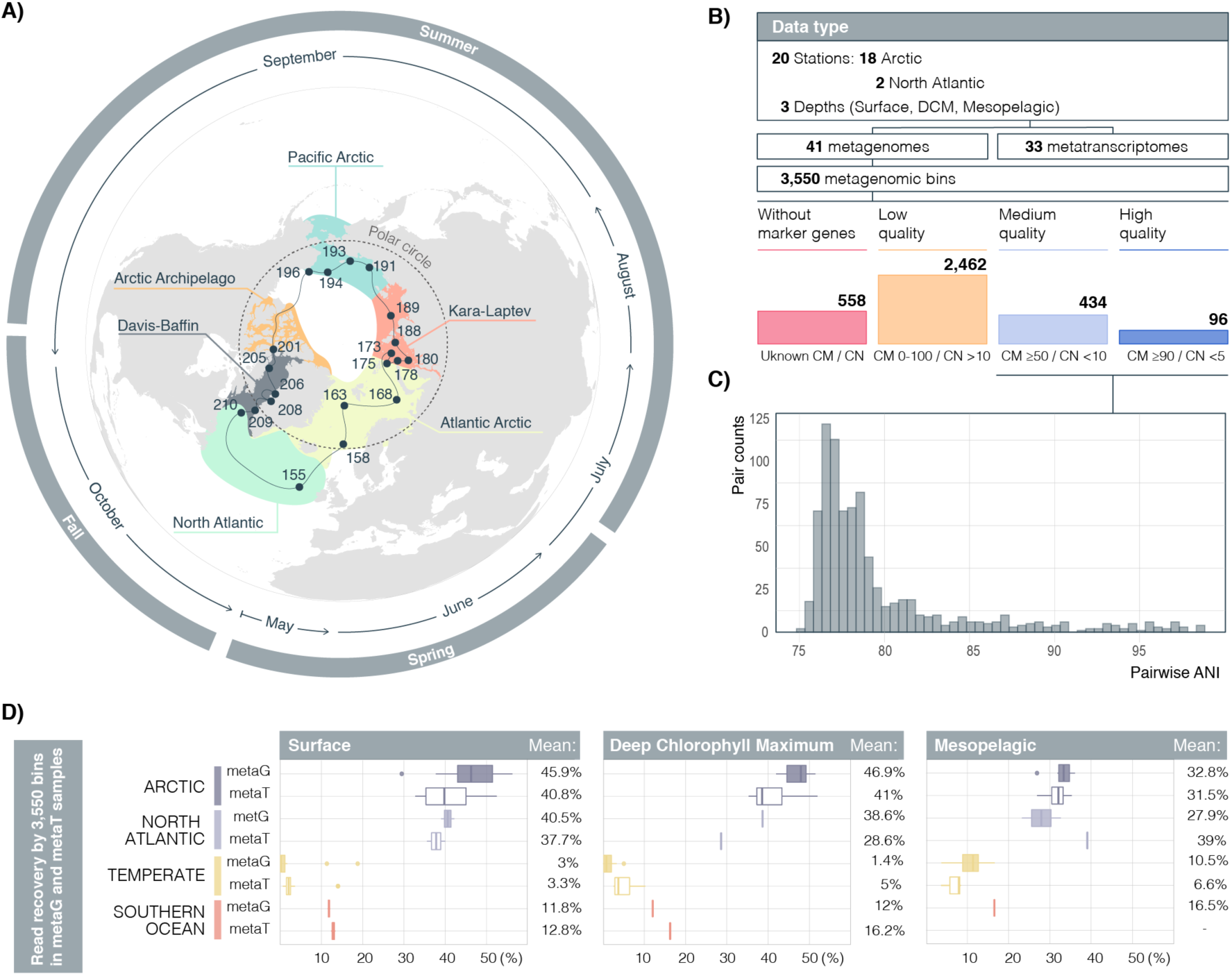
Metagenomic genome reconstruction of *Tara* Oceans Polar Circle expedition. **A)** Ship’s trajectory and the stations from which metagenomes and metatranscriptomes samples are available. Colored areas highlight the sampled regions: five Arctic regions and the sub-Arctic North Atlantic. The Polar Circle (66°N) is shown with a dashed line. Outer circles show the month and season of sampling during the circumnavigation, starting in May 2013. **B)** Outline of the polar metagenomics and metatranscriptomics dataset, the number of bins assembled from metagenomic samples and their quality-based classification, measured by combining genome completeness (CM) and contamination (CN). Only those 530 bins of medium and high quality (MQ and HQ) were denoted as Arctic MAGs. **C)** Pairwise Average Nucleotide Identity (ANI) comparisons of 530 medium quality (MQ) and high quality (HQ) MAGs, showing that only 8 pairs could be considered the same species (ANI >96%). **D)** Distribution of metagenomic (metaG; filled box plots) and metatranscriptomics (metaT; empty box plots) reads’ recovery by all 3550 reconstructed bins per sample. Samples are divided by layer (columns) and latitudinal range (purple boxes for *Tara* Oceans Polar Circle, yellow boxes for temperate samples from *Tara* Oceans Expedition and red boxes for Southern Ocean samples from the *Tara* Oceans Expedition). Mean percentage of read recruitments per group of samples is indicated at the right side of each plot.

This metagenomic genome reconstruction strategy provided 3,550 bins. Their genome based taxonomic classification^39^ resulted in 1,834 bins classified as Bacteria and 146 as Archaea. The remaining unclassified bins (1,570) could have eukaryotic or viral origins, or could not be classified due to a lack of single-copy core genes.

The complete set of 3,550 bins recovered almost half of the dataset of Arctic metagenomic reads (43.3% of Arctic metagenomes and 35.1% of North Atlantic metagenomes, **Figure 1D**). In turn, a subset of 725 Arctic bins that fulfilled the quality standards used by Delmont et al. 2018 (completeness >70% or assembly size > 2Mbp), recovered 23% of Arctic metagenomic reads (**Figure S2**). This is a three-fold difference compared to the 6.84% read recovery by the 892 MAGs generated in Delmont et al. 2018 with *Tara* Oceans metagenomes (which excluded the Arctic sampling)^38^. Our high read recovery could be due to methodological variations (co-assembly and binning strategy, read mapping and filtering) but also to the lower diversity reported in polar prokaryotic communities, compared to those from the temperate ocean^40^.

Interestingly, mean metagenomic recruitments in the Arctic’s mesopelagic were lower than in the photic layer (**Figure 1D**), probably indicating that we are missing genomes from deep Arctic waters. In addition, the mean metagenomic read recovery of Arctic bins increased with depth in temperate and Southern Ocean metagenomes, suggesting that some Arctic genomes may reach the mesopelagic layers of other latitudes through ocean circulation^30,41^. To obtain biogeographic patterns of our Arctic bins at a global scale, we used metagenomes and metatranscriptomes collected from photic and aphotic layers in all the oceanographic regions sampled by the *Tara* Oceans expedition (more details in Materials and Methods). Metatranscriptomic read recruitment in the photic layers of Southern Ocean was four-fold that of temperate samples, suggesting a preference for polar latitudes of certain bins and confirming bipolar expression patterns (**Figure 1D**).

We detected a positive correlation between metagenomic and metatranscriptomic read recruitments by Arctic bins. Similar correlations have been found at the gene level in the eastern subtropical Pacific Ocean^42^, in the global marine microbiome explored by *Tara* Oceans^31^, suggesting that, as may be expected, expression profiles depend on gene abundance^43,44^. The strength of the correlation decreased with depth and was weaker in temperate latitudes (**Figure S3)**, where bins tended to recruit fewer reads from metatranscriptomes than metagenomes. This could be associated with genomes that have been vertically exported from the photic zone to the mesopelagic and/or transported by deep ocean currents to more temperate latitudes. Higher species richness of temperate mesopelagic waters compared to the Arctic could also affect this result^40^. Individual metatranscriptomic recruitments tend to be lower than metagenomic recruitments in temperate latitudes in all layers (**Figure S3**), suggesting that even though the deep currents could connect polar prokaryotes, most cells probably remain in resting stages during transit through non-polar latitudes until reaching favorable habitats in the Southern Ocean^45^. These results reinforce the polar habitat preference of a significant fraction of our *Tara* Arctic genomic dataset.

Following published quality thresholds^36^, the 3,550 bins were classified into four quality groups based on genome completeness and quality values (**Figure 1B, Table S1**): 96 high quality bins (HQ, manually curated, with ≥90% completeness and <5% contamination), 434 medium quality bins (MQ, with ≥50% completeness values and <10% contamination) and 2,642 low quality bins (LQ) bins. Due to a lack of phylogenetic marker genes, 558 bins remained unclassified as their quality could not be estimated. The 530 HQ and MQ bins with sufficient quality ratings were denoted MAGs and are presented in this study as the Arctic MAGs Catalogue. Due to the spatio-temporal and environmental heterogeneity covered by the *Tara* Oceans sampling design, the Arctic MAGs catalogue represents the most comprehensive resource of uncultured prokaryotic genomes from the Arctic Ocean to date.

### DIVERSITY, NOVELTY AND ABUNDANCE OF THE ARCTIC MAGs CATALOGUE

The Arctic MAGs catalogue is composed of a high diversity of non-redundant MAGs. Only eight combinations (0.006%) out of the 140,185 possible genome pairs could be considered to be from closely related species, as they showed Average Nucleotide Identities (ANIs) larger than 96%^46^ (**Figure 1C**). Collectively, our analyses indicate that the Arctic MAGs represent consensus bacterial and archaeal genomes of 526 non-redundant species.

Assembling conserved genes such as the ribosomal operon is a common issue in MAG reconstruction. This is due to the difficulty in resolving *de novo* assembly of short reads using k-mer based methods in a high inter-species sequence identity scenario^47^. In our study, only 27 of the MAGs (5%) contained full or partial ribosomal RNA genes (**Figure S4)**, and therefore, we assessed their taxonomic annotation and novelty through a phylogenomics approach against a database that includes both cultured and uncultured taxa^39^. The Arctic MAGs catalogue included 473 Bacteria and 58 Archaea, assigned to 21 different known phyla (**Figure 2A**), with more than 75% of unclassified bacterial and archaeal genomes at the species level (**Figure 2B**). More details about the taxonomic annotation of Arctic MAGs can be found in the Supplementary Information.

**Figure 2.**
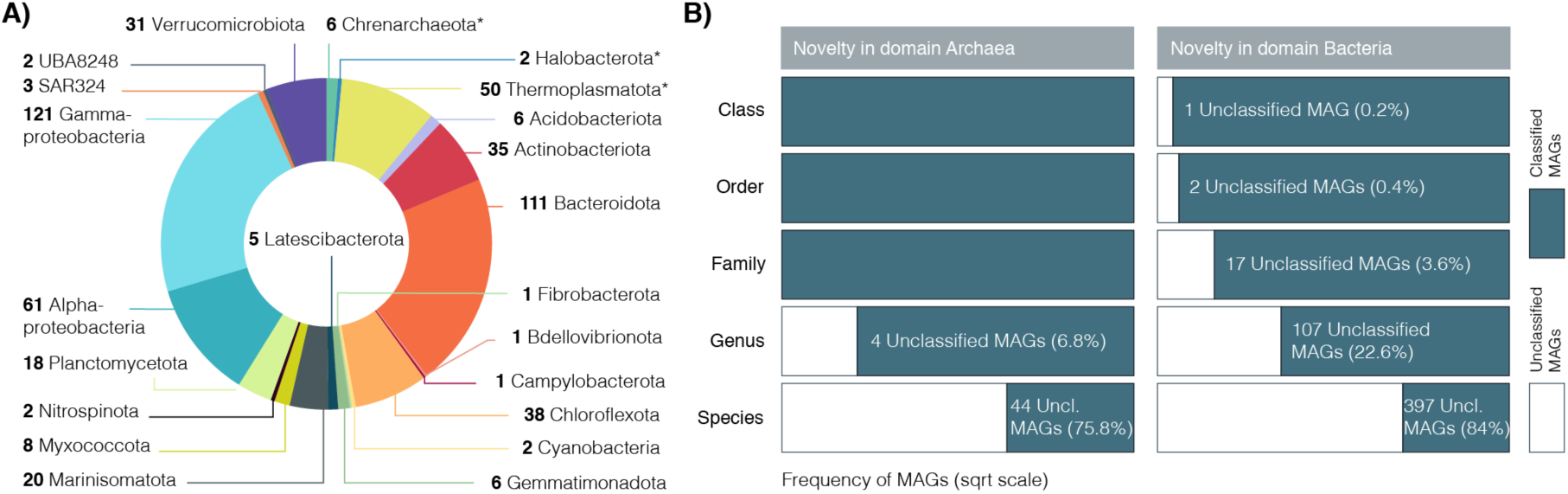
Taxonomical annotation and novelty of Arctic MAGs. **A)** Phylogenomics-based taxonomic classification of the 530 Arctic MAGs dataset at the phylum level (except for Proteobacteria that have been split at the class level). Archaea phyla are highlighted with an asterisk, annotations without asterisk belong to the Bacteria domain. **B)** Stacked barplot for novelty quantification of the Arctic MAGs (X axis) at different taxonomic ranks (Y axis). Taxonomically unclassified portion is depicted in white, taxonomically classified portion is shown in blue. White labelling refers to the unclassified fraction. Frequency of MAGs in axis X is shown in square root scale.

The degree of taxonomic novelty in our dataset is particularly high (**Figure 2B**). Four archaeal MAGs could not be classified beyond the family level, and 44 (75% of the Archaeal MAGs) could not be classified as any known species. In the Bacteria domain, one MAG could not be classified further than phylum Latescibacterota (formerly Latescibacteria), and another could not be classified beyond class Lentisphaeria (phylum Verrucomicrobiota). Novelty increased towards lower taxonomic ranks, with 22% of Bacteria MAGs having an unassigned genus and 84 % of the MAGs belonging to unknown species. Given that most Arctic microbial diversity surveys have relied on PCR-based approaches, using primers designed based on available sequences^20,30,48^, an important fraction of the microbial taxa in this ecosystem may have been consistently missed.

The catalogue contains a majority of rare Arctic taxa (**Figure S5**). The 12 most abundant MAGs, recruiting at least 200 RPKGs (reads per genomic kilobase and sample gigabase) belong to Bacteroidota, Actinobacterota, Alphaproteobacteria, Gammaproteobacteria and the SAR324 phyla. None of them could be classified further than genus, while one of the SAR86 Gammaproteobacteria could not be classified further than family.

A significant positive correlation between whole-genome metagenomic and metatranscriptomic read recruitment in Arctic samples (**Figure S6**) was strong (r>0.7) in 35% of the identified phyla (Crenarchaeota, Bacteroidota, Latescibacterota, Marinisomatota, Planctomycetota, Proteobacteria and Verrucomicrobiota), moderate (r between 0.5-0.7) in 15% (Acidobacteriota, Gemmatimonadota and Actinobacteriota) and weak (r between 0.3-0.5) in 15% of the detected phyla (Thermoplasmatota, Chloroflexota and Myxococcota).

The Arctic MAGs catalogue contains a set of very diverse non-redundant Arctic genomes. Most of them belong to unknown lineages and are representative of both abundant and rare species in Arctic waters, active in terms of gene expression.

### METABOLIC POTENTIAL AND EXPRESSION AMONG ARCTIC MAGs

#### Prevalence of mixotrophy in Arctic prokaryotic genomes

The greenhouse gas CO_2_ is central in the global carbon cycle, and the Arctic Ocean is considered as a sink for atmospheric CO_2_^49,50^. Although primary production in Arctic waters is mainly performed by eukaryotic phytoplankton^23^, inorganic carbon fixation by prokaryotes in the dark might be an important process, particularly during the polar night. For example, nitrification, primarily performed by Crenarchaeota, was detected in deep and surface waters during winter in the Western Arctic^15,16,51^. Similarly, the potential for carbon fixation of certain MAGs from Arctic and Antarctic marine metagenomes was recently reported^33^. However, the relevance and ubiquity of different inorganic carbon fixation pathways across different Arctic regions, depths and seasons is unknown, as is the identity of the potential key players. Although the whole functional annotation is available for the Arctic MAGs (**Supplementary Information**), a selection of 120 marker genes (**Table S2**) representative of carbon fixation processes and energy metabolism was first investigated.

Fifteen Arctic MAGs (2.8% of the total) belonging to seven different phyla contained RuBisCo (KEGG’s K01601, K01602 or both), or RuBisCo and phosphoribulokinase (K00855) and were active in the six studied Arctic regions at all depths (**Figure 3A**). Among them, we report for the first time RuBisCo containing MAGs annotated to the bacterial phyla Latescibacterota and UBA8248 (previously Tectomicrobia). Out of the 15 RuBisCo-containing MAGs, we could retrieve 14 RuBisCo large-chain sequences. These could be classified into phylogenetic groups corresponding to the RuBisCo “Forms” I, II, III-a, IV and IV-like defined in previous studies^52,53^ (**Figure 3B**).

**Figure 3.**
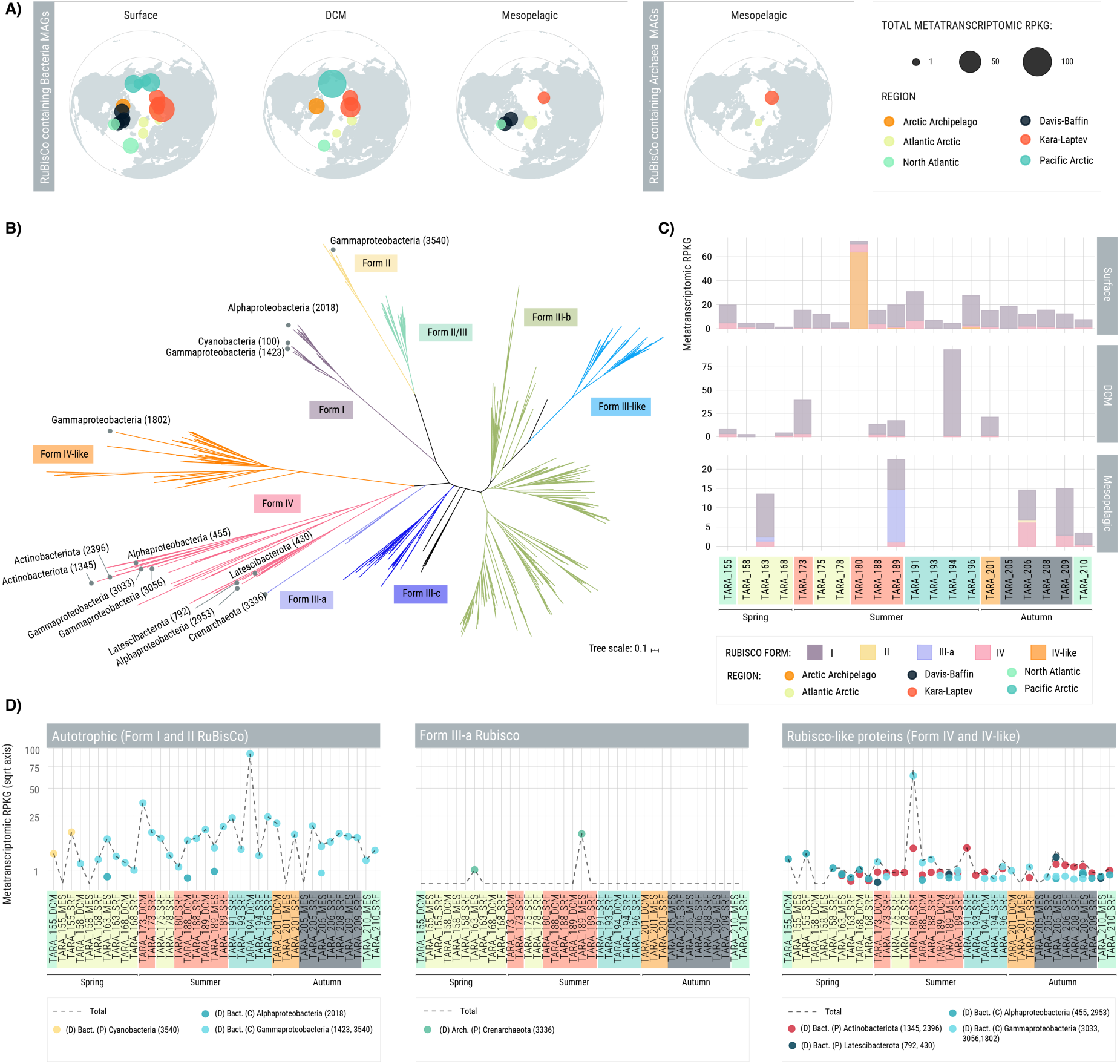
Potential autotrophy in RuBisCo coding MAGs. **A)** Polar maps with the accumulated metatranscriptomic RPKGs of 15 Arctic MAGs encoding at least one subunit of RuBisCo for the Calvin cycle pathway (K01601, K01602), color-coded by Arctic region. The size of the dot is proportional to the accumulated metatranscriptomic RPKGs. **B)** Maximum-Likelihood phylogenetic reconstruction of the 15 RuBisCo large-chain (K01601) aminoacid sequences found in Arctic MAGs, colored by RuBisco form. **C)** Stacked barplot of metatranscriptomic RPKGs recruited by RuBisCo coding MAGs, colored by the RuBisCo form found in their genomes. **D)** Metatranscriptomic RPKGs of RuBisCo containing MAGs collapsed by phylum (or class in the case of Proteobacteria MAGs) and separated by form. Black dashed line represents the total recruited metatranscriptomic RPKGs by RuBisCo coding MAGs in every sample and numbers in parenthesis in legend display the MAG’s identification code.

RuBisCo Forms I and II are directly involved in the autotrophic CO_2_-fixing Calvin-Benson-Bassham (CBB) pathway. Form I was found in one Cyanobacteria MAG (*Synechococcus* sp.), one Alphaproteobacteria (unknown MAG from order UBA2966) and one Gammaproteobacteria (a novel *Thioglobus* sp.). Genetic expression of Form I containing MAGs dominated in all photic samples (surface and DCM) and in five mesopelagic samples regardless of season (**Figure 3C**). Form II was detected in a novel member of the family Thioglobaceae (MAG 3540) expressed only in a mesopelagic sample of the eastern Davis-Baffin (TARA_206). Since activity of the photosynthetic *Synechococcus* was only detected in the TARA_155 North Atlantic samples, expression of MAGs containing RuBisCo Forms I and II in the Arctic suggests a larger contribution of chemoautotrophic processes.

*RuBisCo* Form III-a was detected in a Crenarchaeota (MAG 3336, *UBA57* sp.), which was only active in two mesopelagic spring and summer samples in the Atlantic Arctic and Kara-Laptev seas (**Figure 3C, D**). RuBisCo Form III-a was first reported as being exclusive of methanogenic Archaea and responsible for autotrophic CO_2_ fixation via the CBB pathway in genomes containing phosphoribulokinase (PRK)^53^. Archaea containing RuBisCo Form III-a but lacking PRK, like Arctic MAG 3336, are proposed to be involved in a modified nucleotide scavenging pathway rather than in the fixation of CO_2_^54^. Interestingly, the Arctic Crenarchaeaota’s RuBisCo gene from MAG 3336 is most similar to a Form-IIIa RuBisCo found in a novel clade of deep-sea heterotrophic marine Thaumarchaeota (HMT) MAGs that form a deeply-branched lineage sister with ammonia-oxidizing Archaea^55^. The Arctic MAG 3336 genome size is similar to those of HMT, characterized as ultrasmall (0.6-0.8 Mb), and both are phylogenomically classified as genus UBA57. Even though they do not belong to the same species (pairwise ANI comparisons (between 94.9-95.1%), these results suggest that they likely share a similar heterotrophic lifestyle.

RuBisco Form IV and IV-like RuBisCo (or RuBisCo like proteins, RLP) do not perform CO_2_ fixation and may be involved in methionine salvage, sulfur metabolism and D-apiose catabolism^56,57^. RLPs were found in two Gammaproteobacteria MAGs (novel *BACL14* sp. and *UBA4575* sp.), two Alphaproteobacteria (*HIMB11* sp. and an unknown Magnetovibrionaceae MAG), two Actinobacteriota (*UBA4592* sp. and a novel *Planktophila* sp.) and two Latescibacterota (unknown MAGs from families GCA-002724215 and UBA2968). Their metatranscriptomic read recruitment, albeit generally smaller than for MAGs with CBB cycle potential, occurs in all samples, while Form IV containing MAGs are expressed both in the photic and mesopelagic layers, Form IV-like containing MAG is only active in the surface of three summer samples (**Figure 3C**).

These results indicate that at least 28% of RuBisCo containing MAGs are potential autotrophs (forms I and II), prevalent in the Arctic Ocean and expressed across all regions, depths and seasons. Nevertheless, RuBisCo containing MAGs possessed multiple (from 15 to 134) protein domains annotated as ATP-binding cassette (ABC) transporters, in charge of sugar, amino acid and oligopeptide transport (**Table S3**) reflecting a likely mixotrophic lifestyle (Crenarchaeota MAG 3336, a potential heterotroph). Mixotrophy is also suggested for the photosynthetic *Synecochoccus* MAG, a lifestyle that has already been reported in other marine Cyanobacteria^58^.

Mixotrophy has also been proposed to be relevant for specific Arctic heterotrophs, which can incorporate CO_2_ in the dark without any net carbon assimilation in processes linked to fatty acid biosynthesis, anaplerotic reactions^13^ or CO-oxidation^59^. The oxidation of carbon monoxide is suggested to serve as a supplemental energy source during organic carbon starvation^59^. This process is catalyzed by carbon monoxide dehydrogenase (CODH; cox genes)^60,61^ and has been found previously in Actinobacterota, Proteobacteria, and taxa from Bacteroidetota and Chloroflexota^59,62^. We found a total of 332 (62%) MAGs containing the coxL (K03520) gene, key for CO-oxidation, to be metabolically active at the time of sampling. These belonged to 10 Bacteria phyla and 2 Archaea phyla and were ubiquitously expressed (**Figure S7**). To our knowledge, this is the first approach to assessing CO oxidation potential by prokaryotes in the Arctic Ocean.

Some key markers for the 3-Hydroxypropionate bicycle (from now on 3-HP, **Figure S8**) and the 3-Hydroxypropionate/4-Hydroxybutyrate Cycle (from now on 3-HP/4-HB, **Figure S9**) were also detected but considering the lack of complementary genes for carbon fixation, such as acetyl-CoA carboxylase in the 3-HP, the autotrophic capacity of these MAGs remains putative, which calls for further metabolic network reconstruction studies.

### Chemolithoautotrophic potential of Arctic prokaryotic genomes

We investigated metabolisms associated with ammonia and nitrite oxidation (**Figure 4**), resulting in five MAGs that contain reliable markers for chemolithoautotrophic processes. We could describe three ammonia-oxidizing archaea (AOA), two annotated as novel *Nitrosopelagicus* spp. (containing 31% of 3-HP/4-HB KEGG module, the full urease complex, and in MAG 1708, the ammonia-monooxygenase coding *amoA*) and one *Nitrosopumilus* sp. (containing 21% of the 3-HP/4-HB KEGG module and the full urease complex). One Alphaproteobacteria (*GCA-2728255* sp.) was classified as an ammonia oxidizing bacterium (AOB) and contained the characteristic nitrification marker hydroxylamine oxidoreductase *hao*. One *Nitrospinia* species *LS-NOB* sp. was classified as a nitrite oxidizing bacteria (NOB), with 35% of reverse TCA cycle module completeness, including ATP citrate lyase, the nitrite oxidoreductase *nxr* and a nitrate/nitrite transporter. Their expression patterns showed an overall preference for mesopelagic depths, especially in the North Atlantic-influenced Arctic stations (in spring and autumn). AOA and AOB are also active in the North-Atlantic influenced mesopelagic of one station of the Kara-Laptev sea, the former recruiting more metatranscriptomic RPKGs, in coherence with previous results^27^. NOB expression is restricted to the photic samples of the North Atlantic spring station TARA_155 and the mesopelagic of the North Atlantic autumn station TARA_210. Bulk nitrite oxidation in euphotic North Atlantic subpolar regions during spring and autumn was described recently ^63^. Nevertheless, to our knowledge, the *LS-NOB* sp. MAG is the first individual NOB representative found to be active in the region in both photic and aphotic layers.

**Figure 4.**
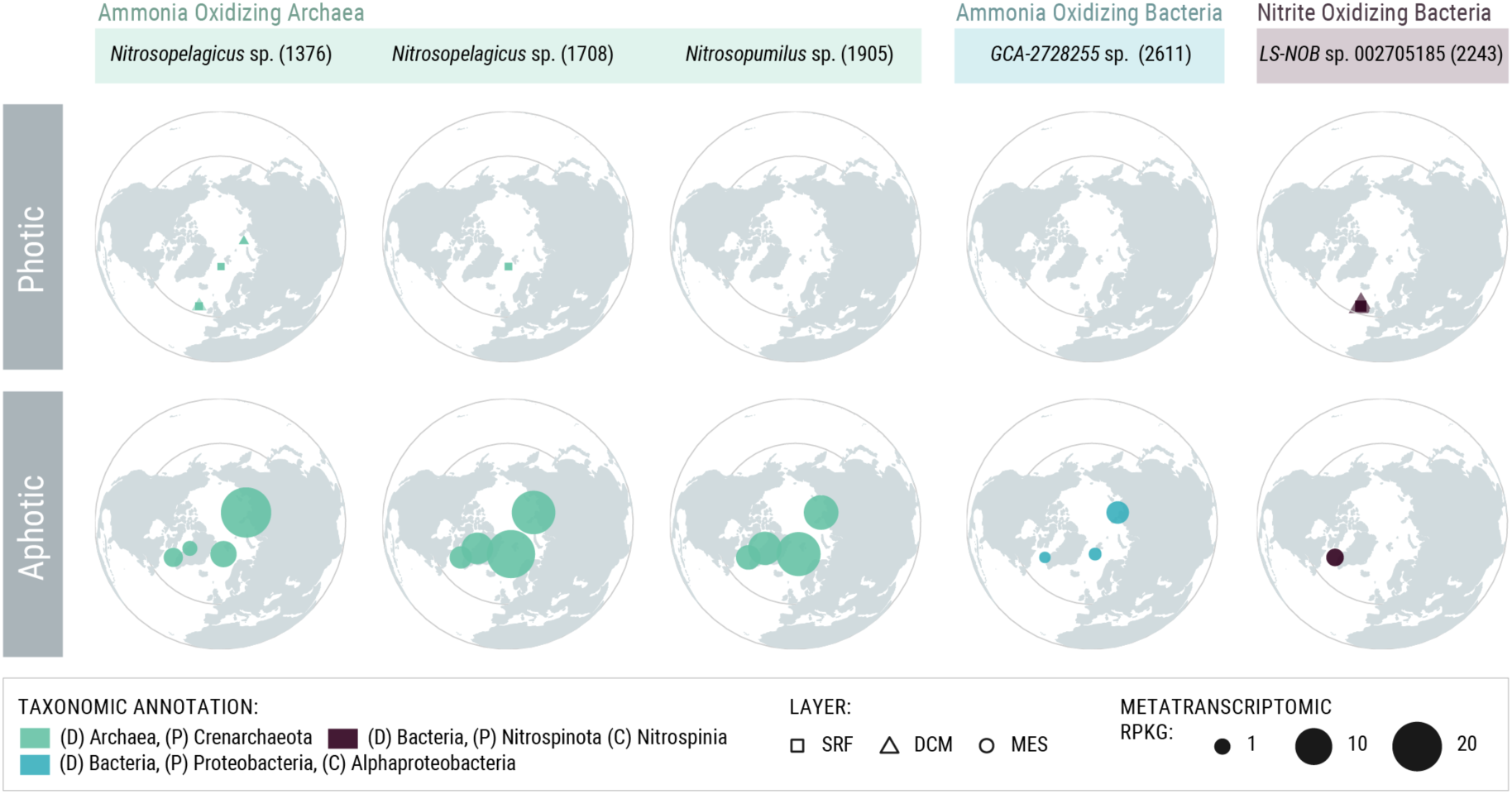
Chemolithoautotrophic Arctic MAGs. Five MAGs contained the specific markers genes to be putative chemolithoautotrophs in the Arctic Ocean. They are classified into Ammonia Oxidizing Archaea and Bacteria and Nitrite Oxidizing Bacteria. Their metatranscriptomic recruitments in RPKGs in depicted by the size of the dots, while their shape indicates the water column layer. Color of the dots depends on the taxonomic annotation of each MAG. Numbers between parenthesis correspond to the MAG’s code.

It therefore appears that the set of Arctic MAGs is made by a majority of heterotrophic and mixotrophic organisms, with a few chemolithoautotrophs that are mostly expressed in the mesopelagic during spring and autumn. Future experimental validation is required to confirm quantitatively the relevance of these processes.

### ECOLOGICAL PREFERENCES AND BIOGEOGRAPHIC PATTERNS

The Arctic MAGs were used as reference genomes in the mapping of metagenomic reads from 68 samples, covering five Arctic regions, the sub-Arctic North Atlantic and representing all the temperate and Southern Ocean oceanographic regions sampled by *Tara* Oceans. Ordination of samples based on Bray-Curtis dissimilarities of MAG composition clearly grouped polar samples together (Arctic and Southern Ocean), separated from temperate samples (**Figure 5A**). This pattern was similar to the clustering of the global *Tara* Oceans samples based on their 16S miTAG profiles (**Figure S10**). This suggests that polar waters contain a unique diversity of prokaryotes, different from temperate regions, and confirms the presence of bipolar taxa, previously described in surveys based on PCR amplicon sequencing^30,64^. Within polar and non-polar samples, MAG assemblages were significantly structured by depth (NMDS with 100 iterations and 0.8 stress value, Permutational MANOVA R^2^=0.138, p-value <0.001) (**Figure 5A**) in agreement with previous studies on the vertical stratification of marine microbial communities^22,30,65,66^.

**Figure 5.**
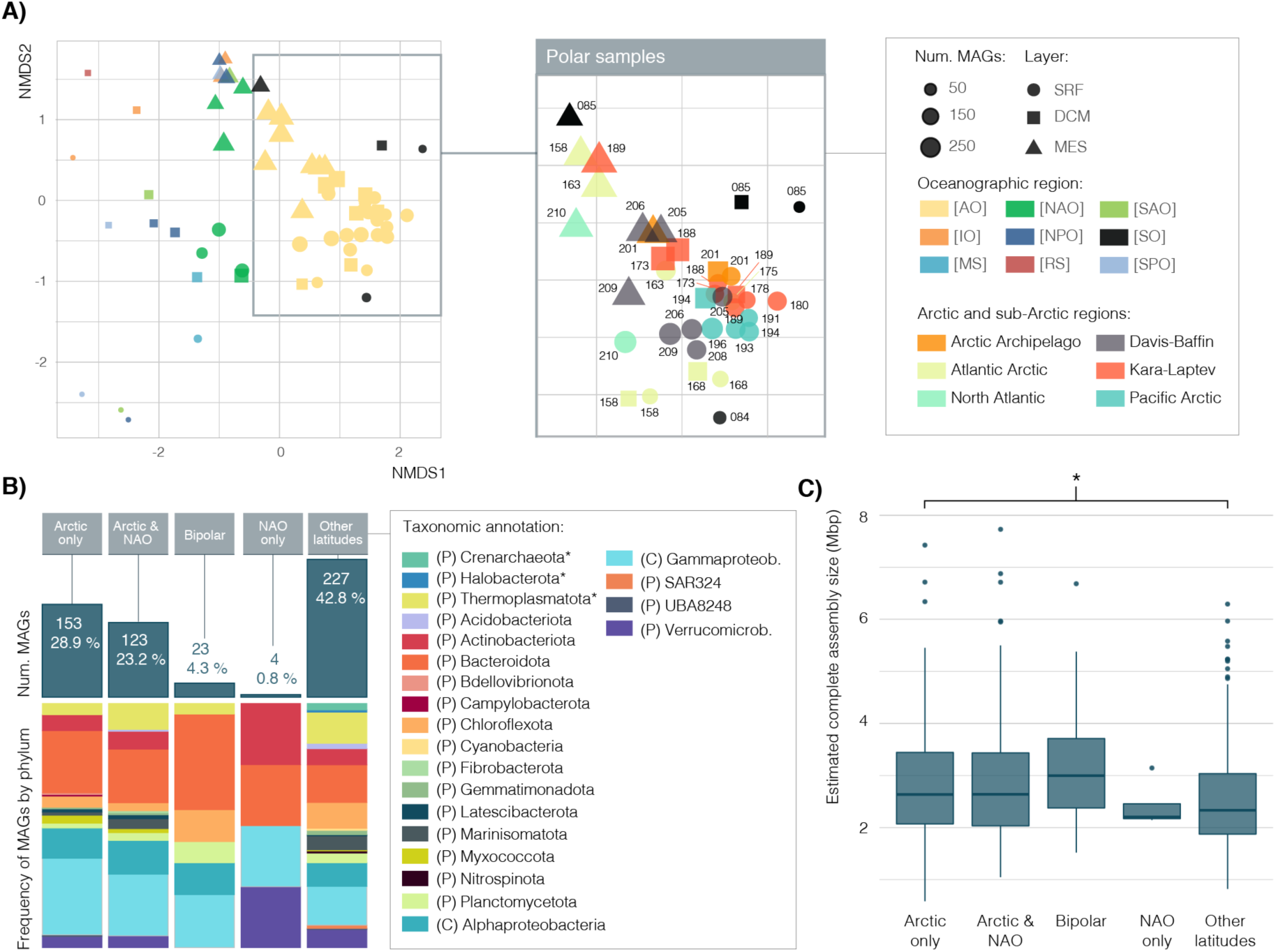
Composition and biogeography of the 530 Arctic microbial MAGs. **A)** Non-metric Multidimensional Scaling (NMDS) ordination of metagenomic samples based on their composition of 530 Arctic MAGs. Shape defines the sample’s layer in the water column (SRF: surface, DCM: Deep Chlorophyll Maximum, MES: Mesopelagic) and the dot size represents the MAG richness (i.e., number of different MAGs) of the sample. The plot on the left shows all non-polar and polar (inside the square) samples, colored by oceanographic region. Oceanographic regions are: Artic Ocean [AO], Indian Ocean [IO], Mediterranean Sea [MS], North Atlantic Ocean [NAO], North Pacific Ocean [NPO], Red Sea [RS], South Atlantic Ocean [SAO], Southern Ocean [SO], South Pacific Ocean [SPO]. The middle and the right panels represent the same NMDS ordination of only polar samples color-coded by season or Arctic/sub-Arctic region, respectively, and labeled with their Station number. **B)** Biogeographic categorization of the 530 Arctic MAGs. Stacked bar plots represent the number of MAGs in each category, colored by taxonomic annotation, and top bars represent the percentage within the medium and high-quality Arctic MAGs dataset. **C)** Between different biogeographic categories. DTK test shows significant differences (p-value < 0.05) between MAGs specific from the Arctic and MAGs present in lower latitudes.

We also delineated the geographic distribution of individual MAGs using metagenomes representative of the global ocean and an astringent read mapping filtering of at least 20% of horizontal genome coverage. We found 153 MAGs (28.9%) present exclusively in Arctic metagenomes, and 23 (4%) showing a bipolar distribution (i.e., recruiting reads only from Arctic and Southern Ocean metagenomes) (**Figure 5B**). A previous study that used samples from ICoMM reported bipolarity of 15% of the generated OTUs^30^. Such a difference with our results might be explained by our genome-centric approach, which might be more conservative than the definition of biogeographic patterns based solely on the 16S rRNA gene and/or the fact that the MAGs only represent a fraction of diversity in these communities. The bipolar subset of MAGs was less rich in prokaryotic phyla diversity than other biogeographic categories, consistent with latitudinal diversity gradients^40^, and lacked MAGs representative of Actinobacteriota and Verrucomicrobiota found in every other studied latitude (**Figure 5B**).

Almost 25% of MAGs were present only in both Arctic and sub-Arctic North Atlantic sample sets. Together with the 29% of Arctic specific and 4% bipolar MAGs, almost 60% of our MAG dataset is represented by polar genomes (**Figure 5B**). When comparing estimated complete genome sizes of the Arctic MAGs, we found that those genomes with an Arctic-only distribution were estimated to be significantly larger (2.9 Mbp on average) than those with a presence in temperate latitudes (2.5 Mbp) (DTK, Dunnett’s Modified Tukey-Kramer Pairwise Multiple Comparison Test p-value <0.05) (**Figure 5C**). However, we did not find significant differences between their coding densities (i.e., the fraction of the MAG annotated as coding sequences or CDS).

In summary, about 30% of our MAGs dataset is exclusively present in Arctic regions and the large genome of Arctic-only MAGs may confer a higher functional and metabolic versatility in the extreme Arctic Ocean environment.

### DISENTANGLING GENERALIST AND SPECIALIST ARCTIC MAGs

To explore which Arctic MAGs display a panarctic distribution or a more restricted distribution, we defined two subsets of genomes based on their niche breadth. On the one hand, we consider the habitat generalists, evenly distributed in the majority of Arctic samples; and on the other hand, the habitat specialists, with an uneven distribution, usually peaking in abundance in a reduced number of samples^67,68^. The latter are thought to be more sensitive to changes in environmental conditions^69–71^, as they might have narrow environmental requirements. Generalists, on the other hand, are less dependent on environmental conditions, have a wide habitat tolerance, and high functional plasticity^72^. In the current scenario of climate change, it is essential to identify which Arctic species may be more susceptible to environmental change.

We calculated the niche breadth of individual MAGs based on their abundance and occurrence across the Arctic metagenomic dataset. For this analysis, each Arctic sample was considered as an individual habitat, as their geographical location, depth in the water column, and season of sampling differed. In line with previous studies^71,73^, the majority of Arctic MAGs (71%) could not be categorized into generalists or specialists, while 21% (n=111) were habitat specialists and 7% (n=38) were generalists (**Figure 6A**). High contributions of specialists have been reported in other polar environmental extremes such as coastal Antarctic lakes^70^ or in highly productive marine sites compared to oligotrophic open ocean stations^74^. Both generalist and specialist MAGs show a similar range in their mean abundances, which contrasts with lower abundances of specialists in niche breadth analyses of Arctic eukaryotes (Karp-Boss et al., submitted).

**Figure 6.**
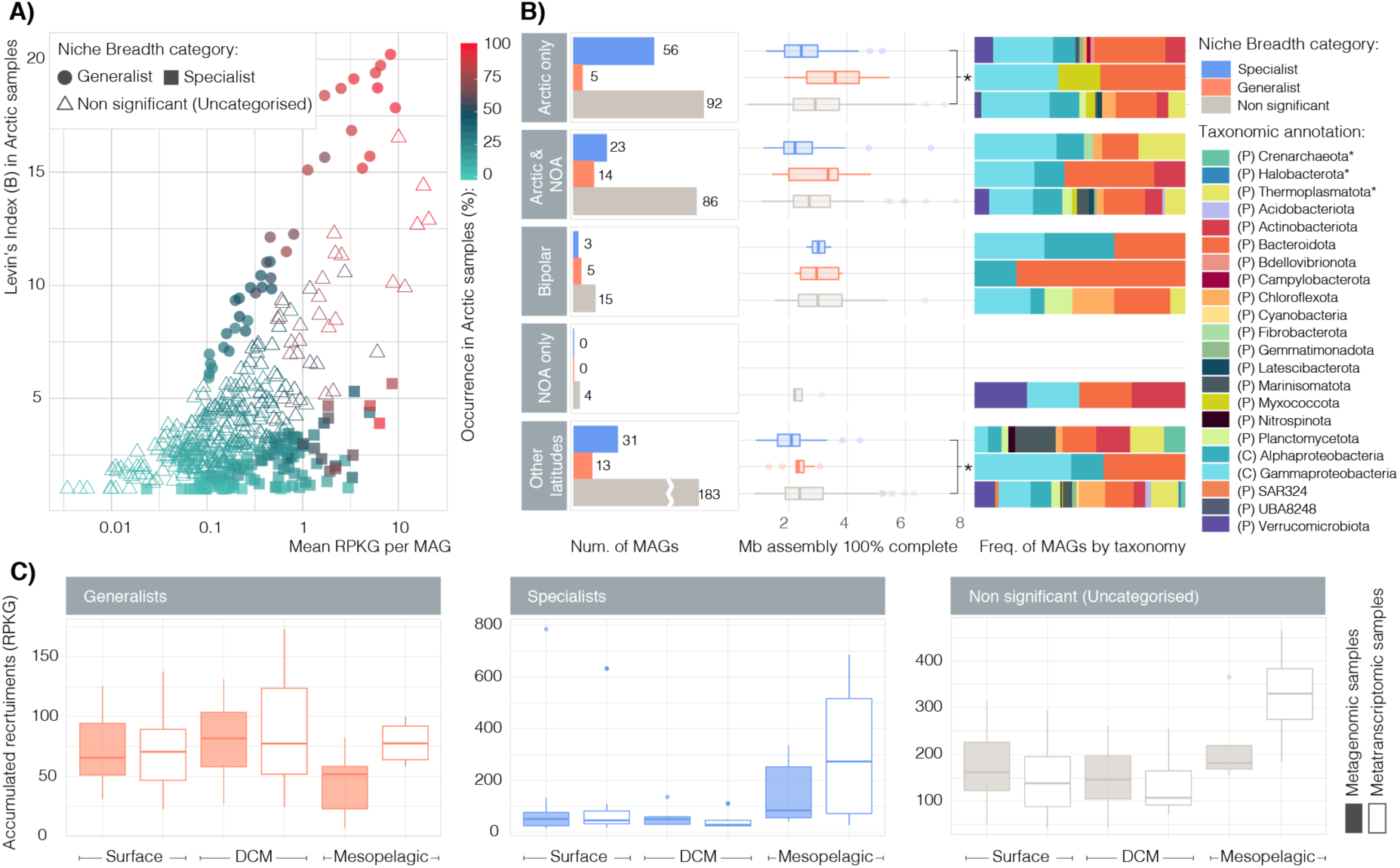
Disentangling generalists and specialists within the 530 Arctic MAGs. **A)** Distribution of Arctic MAGs based on their mean read recruitments in Arctic metagenomic samples (RPKG, X axis) and their Levin’s Index (i.e, niche breadth, Y axis). The color gradient depicts the occurrence (i.e., % of samples where a given MAG is present) in the Arctic metagenomic dataset and shape indicates their niche breadth category (generalists, specialists and uncategorised). **B)** Number of habitat generalists (orange), specialists (blue) and uncategorised MAGs (grey) in each biogeographic category shown in bar plots. The adjacent boxplots show the distribution of assembly sizes within each subcategory (upscaled to 100% of genome completeness) and statistically significant differences have been highlighted with an asterisk (DTK test p-value <0.05). Stacked barplots indicate their taxonomic composition at the phylum level. Asterisks in the taxonomic annotation legend indicate phyla from domain Archaea, lack of asterisk indicates domain Bacteria. **C)** Abundances of generalists (orange), specialists (blue) and uncategorised (grey) MAGs in metagenomic (filled boxplots) and metatranscriptomic (empty boxplots) samples across the three ocean layers. There are no significant differences between the groups.

As habitat generalists are likely to adapt to a broader range of habitats due to their functional plasticity^72^, we expected their estimated complete genome size to be larger than that of habitat specialists. This difference was apparent but not statistically significant in the median genome size of MAGs that showed Arctic and North Atlantic distributions (DTK, Dunnett’s Modified Tukey-Kramer Pairwise Multiple Comparison Test) (**Figure 6B**). Overall, specialist MAG genome size was significantly lower than those of uncategorized MAGs (DTK test p-value <0.05) (**Figure S11**).

While generalists were assigned to Bacteria phyla Actinobacterota, Proteobacteria, Bacteroidetota and Myxococcota (**Figure 6B**), specialists displayed a larger taxonomic diversity, including the archaeal phyla Thermoplasmatota and Crenarchaeota.

Interestingly, the number of metagenomic and metatranscriptomic RPKGs belonging to specialist and generalist MAGs was similar in the photic zone. In contrast, mesopelagic samples had a higher proportion of metagenomic and metatranscriptomic RPKGs belonging to specialist MAGs (**Figure 6C**). This difference might be explained by nutrient availability and niche compartmentalization in the deeper waters, like different composition and labile stage of euphotic zone-derived sinking particles, and/or buoyant particles that are produced autochthonously at depth^75^, in contrast with the wider gradients in nitrate, temperature and salinity of the upper Arctic Ocean (**Table S4**). Since the abundance or expression of microbial generalists or specialists could not be explicitly linked to any of the environmental variables tested (**Figure S12**), it is likely that community turnover in polar communities, suggested to drive changes in the community’s gene expression in response to ocean warming, could also transcend niche breadth.

### POLAR OCEAN PROKARYOTIC SENTINELS

In order to define key prokaryotic genomes specific to polar regions, and whose existence may be threatened by the expected changes in the polar environment (i.e., sentinel genomes), we examined MAGs that displayed an exclusively polar (either Arctic or bipolar) distribution and were most expressed in every sample and within their niche preference group (specialist, generalist, uncategorized). A total of 62 MAGs were selected based on these criteria **(Figure 7)**. These sentinel MAGs (7 generalists, 25 specialists and 30 uncategorized) may represent potential ecologically relevant taxa in the polar ecosystem that we advocate to monitor as a means to assess the health status of the Arctic Ocean.

**Figure 7:**
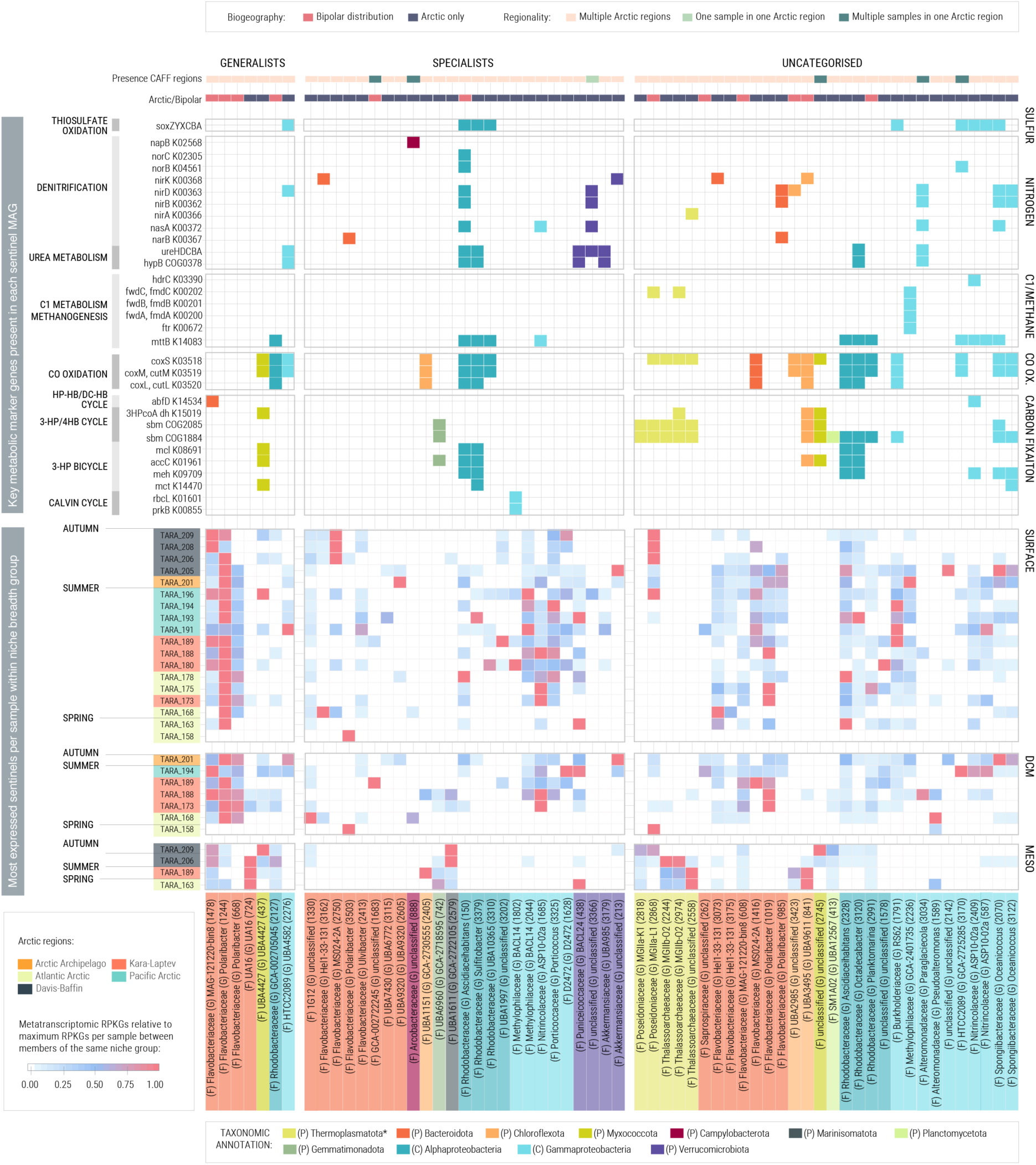
Expression patterns and metabolic potential of sentinel polar MAGs in the Arctic Ocean. The plot contains a selection of 62 MAGs, which are the most expressed per sample within the subset of polar specific MAGs that are either generalists, specialists or uncategorized. The top tile plot represents which of the selected marker genes are encoded in each of these MAGs. The bottom heatmap represents the relative expression of each of these MAGs (X axis) in each sample (Y axis). Recruitment normalizations were done for every niche breadth category. Samples on the Y axis are colored based on the Arctic region they belong, and the sampling season is indicated. MAGs on the X axis are colored based on phylum and the number in parenthesis corresponds to the identification number of each MAG.

As an example, *Polaribacter* is one of the most common genera in polar waters and its bipolar distribution had been described previously^76^. *Polaribacter* spp. was one of the genera that showed the highest number of MAGs with a bipolar distribution and highest expression across most of the photic samples (**Figure 7**). *Polaribacter* MAGs were assigned to both generalists and specialists. Interestingly, in a parallel study using the same Arctic samples, *Polaribacter* is predicted to be an ecologically central species within a cross-domain interactome community enriched in polar stations, being identified as one of the most connected taxa (Chaffron et al, submitted) and ranking high according to the General keystone index (Keystone Index Rank = 8 out of 1498)^77^. Another heterotrophic Flavobacteria (UA16 family) dominated gene expression in the mesopelagic, together with a MAG from the Myxococcota family UBA4427 (**Figure 7**). Photic generalists were mostly heterotrophic but we also found a generalist annotated as Myxococcota thriving in the mesopelagic with a putative autotrophic metabolism.

The expression patterns of polar sentinel specialists were not dominated by any particular taxonomic group (**Figure 7**). We found potential for autotrophic metabolism through the Calvin cycle in a MAG related to Gammaproteobacteria (Methylophilaceae) and the 3-hydroxypropionate cycle in a novel Gemmatimonadetes species and two Alphaproteobacteria (Rhodobacteraceae). The majority of highest metatranscriptome recruitments in photic summer samples by sentinel specialists were associated with Gammaproteobacteria and Alphaproteobacteria. In contrast, spring and autumn photic samples showed highest expression values from *Polaribacter* and other Flavobacteria and Verrucomicrobia. Most of these Gammaproteobacteria were heterotrophic with denitrifying potential. In the mesopelagic, the most active specialists in terms of gene expression were MAGs from the phylum Verrucomicrobiota in spring, Chloroflexi in summer and Marinisomatota in autumn.

For the MAGs that did not fall into a niche breadth category, we found that higher expression in surface waters through spring and summer was associated with; i) those showing heterotrophic metabolism, e.g., MAGs from a novel Alphaproteobacteria family (MAG 2142) and Bacteroidota, and ii) potentially chemolithoautotrophic or mixotrophic MAGs, belonging to Chloroflexi and Alpha- and Gammaproteobacteria. During this time, Thalassoarchaeaceae MGGIIb and Chloroflexota were the most active in the mesopelagic samples. Autumn photic samples were clearly dominated by Archaea MAGs from the family Poseidoniaceae MGIIa. *Oceanicoccus* Gammaproteobacteria were predominantly active in the DCM and a novel family of Planctomycetota was most expressed in the mesopelagic (**Figure 7**).

Overall, we uncovered a pool of 62 Arctic sentinel MAGs (11% of the Tara Arctic MAG dataset), of which seven were classified as habitat generalists and 25 as habitat specialists. They were present only in polar latitudes and were highly active in the Arctic Ocean. While sentinel generalists seem to be mostly heterotrophic, sentinel specialists display a wider variety of metabolic markers, including autotrophic potential and denitrifying genes. Sentinel specialists appear to be candidates for colonization of a wider variety of niches to exploit the available chemical energy. However, their reduced niche breadth would make them vulnerable to the effects of climate change in the Arctic waters.

## Conclusions

The assembly of a unique catalog of 530 Arctic metagenomic assembled genomes (MAGs), of which more than 75% represent novel species, highlights that we have not yet discovered either the full taxonomic nor the functional diversity of microbial communities in the Arctic Ocean. This study also suggests a new approach for the generation of metagenomic bins based on the co-assembly of pooled metagenomes more similar in their community composition and later binning of all the contigs together, which has resulted in a bin dataset that recovers half of the sequenced community and whose MAGs show very low redundancy. Our in-depth genome-centric analysis of novel lineages in the Arctic Ocean highlights those with the highest genetic expression, thus more likely to be active in Arctic seawaters, and provides a high number of new Arctic reference genomes including thousands of potentially non-prokaryotic bins for the exploration of keystone Arctic viral or eukaryotic genomes.

The eco-genomics perspective presented here enabled us to define the prevalence of mixotrophic activity and chemolithoautotrophy from spring to autumn and to identify potential sentinel species in the Arctic’s seawater ecosystem, some of which might be more susceptible to the effects of climate change due to their restricted niche breadth. The description of their functional capabilities and relevance in terms of genome expression is also key for future design of monitoring, experiments and ecosystem models in this rapidly changing environment.

## Materials and methods

### Sample and environmental data collection

As described previously^31^ genetic and environmental data were collected during the *Tara* Oceans expedition (2009-2013), which includes the *Tara* Oceans Polar Expedition (TOPC, 2013). Polar stations had absolute latitudes above 64°. Sampling was conducted within the epipelagic (surface / SRF, 5-10 m and deep chlorophyll maximum / DCM, 20-200 m) and mesopelagic layer (MES, 20-200 m). The sampling strategy and methodology have been described elsewhere^35^. Environmental data measured or inferred at the depth of sampling are published at the PANGAEA database (https://doi.org/10.1594/PANGAEA.875582).

### Extraction and sequencing of DNA and cDNA

Metagenomic DNA and RNA were extracted from prokaryote-enriched size fraction filters (0.2 um-3um) as previously described^78^. A detailed description of the DNA sequencing protocols is given in^31^.

### Co-assembly, binning and curation

#### Co-assembly

Bins were generated from 41 Tara Oceans Arctic metagenomes, including 28 samples from the photic layer (20 from the surface and eight from the DCM), nine from the mesopelagic layer and four integrated samples, in which waters from different layer were mixed. In order to maximize the recovery of environmental genomes from the dataset, we opted for an approach that involved the co-assembly of several samples together, hence increasing the sequencing depth for each co-assembly while keeping the computational needs attainable. The pools of samples to be co-assembled were chosen based on their taxonomic composition. Samples that clustered together in an NMDS based on 16S miTag abundance profiles were assembled jointly with megahit (v1.1.2, --presets meta-large --min-contig-len 2000; **Table S5**; **Figure S13**)^79^. All assembled contigs were pooled together and de-replicated with cd-hit-est v4.6.8-2017-0621, compiled from source with MAX_SEQ=10000000, options -c 0.99 -T 64 -M 290000 -n 10^80^, reducing the dataset from 3.95 M to 1.91 M contigs.

#### Binning and curation

The reads of the input metagenomes reads were back-mapped to the remaining contigs with bowtie2 v2.3.2^81^ with default options, keeping only mapping hits with quality larger than 10 (samtools v1.5; options -q 10 -F 4)^82,83^. Mapping hits were processed with jgi_summarize_bam_contig_depths from metabat2 v2.12.1^83^ with options --minContigLength 2000 --minContigDepth 1 and then binned with metabat2 with default options.

The completeness and contamination of each bin, as well as a first estimation of their taxonomic classification, based on single-copy marker genes was assessed with checkM v1.0.11 ^84^ using the lineage_wf workflow.

Contigs of 96 bins with estimated completeness larger than 95% and contamination lower than 5% were reassembled in Geneious v10.2.4 with minimum overlap identity 95%, maximum mismatches per read 5, no minimum overlap and with no gap allowed options to find overlaps that allowed to reduce the genome fragmentation, and the results were curated manually. These were considered to be high quality (HQ) MAGs. Additionally, contigs of 434 bins with estimated genome completeness larger than 50% and contamination lower than 10% were also re-assembled with cap3 v021015^85^ with overlap length and percent identity cut-offs of 25 bp and 95% respectively. These were considered to be medium quality (MQ) MAGs.

All 3,550 genomes were given a numeric identifier, with the prefix “TOA-bin-”, that stands for *Tara* Oceans Arctic bin.

### Taxonomic and functional annotation

All 3,550 bins were classified taxonomically with GTDBTk v0.3.2^86^ using the classify_wf workflow. Genome completeness and contamination estimates were reassessed with checkM as above. For those bins encoding the 16S rRNA gene, their taxonomic annotation was done using SILVA 132 database and SINA aligner tool v1.2.11 with a minimum of 50% of identity (higher thresholds could not classify the ribosomal genes) and Last Common Ancestor algorithm. (**Table S6**).

Functional annotation of 530 MAGs, including gene prediction, tRNA, rRNA and CRISPR detection was done with prokka v1.13^87^ using default options and the estimated Domain classification from checkM as the argument in the --kingdom option. Additionally, predicted coding sequences were annotated against the KEGG orthology database (KEGG release 2019-02-11)^88^ with diamond v0.9.22^89^ using options blastp -e 0.1 --sensitive, and against the PFAM database release 31.0 using hmmer v3.1b2^90^ and options --domtblout -E 0.1. Functional annotation of MAGs can be accessed in the Supplementary Information.

### Genome redundancy analysis

Average nucleotide identity (ANI) was calculated with fastANI v1.2 and default options^91^ was estimated for each possible pair of MAGs with more than 50% of genome completeness and less than 10% of genome contamination to check whether the reconstructed genomes could belong to the same species (defined at >95% ANI). As alignment fraction between genomes lower than 20% may provide spurious large ANIs, the average amino acid identity (AAI), which considers only the fraction of orthologous genes, was also estimated (compareM v0.0.23 with default options; https://github.com/dparks1134/CompareM).

### Read recruitments

#### Selection and subsampling of samples

The samples chosen for read recruitment include the 37 surface, DCM and mesopelagic metagenomes from *Tara* Oceans Arctic Stations (**Figure 1A and 1B**), the four *Tara* Oceans metagenomes sampled in the Southern Ocean and a selection of 27 *Tara* Oceans expedition metagenomes from temperate latitudes (**Table S4, Figure S13**). These were selected based on their sequencing depth (that had to be at least as large as the smallest *Tara* Oceans Arctic metagenome), geographic location (covering the different oceans and seas sampled by the *Tara* Oceans expedition), and the coverage of different water layers. For the metagenomic samples selected, recruitments were also done with their available metatranscriptomes (33 from the *Tara* Oceans Arctic Stations, three from the Southern Ocean and 17 from the temperate ocean). Paired-end libraries were used individually for Fragment Recruitment Analysis after cleaning and a step of random subsampling. The latter was done with DOE JGI’s BBTools’ reformat.sh script v38.08 (https://sourceforge.net/projects/bbmap/), selecting as subsampling value the smallest sequencing depth of the *Tara* Oceans Arctic expedition meta-omic dataset (i.e 140,658,260 and 45,212,614 fragments for metagenomic and metatranscriptomic libraries respectively). Read length was 101 bp.

#### Competitive fragment recruitment analysis

Nucleotide-Nucleotide BLAST v2.7.1+ was used to recruit metagenomic and metatranscriptomic reads similar to any of the 3,550 Arctic bins. Blast is slower than other high-throughput (HT) aligners but allows for finer-tuned alignment parameters, plus it is the gold standard against which all HT aligners are compared. Recruitment was competitive, meaning that individual samples were aligned against the pooled contigs of all 3,550 bins. Blast alignment parameters were the following: -perc_identity 70, -evalue 0.0001. Only those reads with more than 90% coverage and mapping at identities equal to or higher than 95% were considered to be representative of the bin. In case of hits with the same e-value, larger bit-score or larger alignment length were used sequentially to choose the best hit. If ties persisted, the best hit was selected at random from the candidate reads. Best hits that corresponded to rRNAs (according to the prokka annotation) were also discarded.

#### Detection and filtering of false-positive recruitments

Putative false positive recruitments were detected and excluded considering their horizontal genomic coverage which was calculated using the R package GenomicRanges^92^.

A minimum horizontal genomic coverage threshold was set testing the effect of different thresholds on the final number of bins recruiting (richness) and the number of samples in which they recruited (occurrence). The variation of species richness in each metagenome was tested for a range of increasing minimum horizontal genomic coverage thresholds (0, 0.1, 0.5, 1, 2, 3, 4, 5, 10, 15, 20, 25, 30, 35, 40, 45, 50, 55, 60, 65, 70, 75, 80, 85, 90, 95, 98, 100). Recruitments in which the horizontal coverage was equal to or higher than the thresholds, were considered true and those covering a smaller percentage of their genome than the cut-off value were discarded.

The number of species present in each metagenome decreased with the increase of minimum horizontal coverage, reaching an apparent saturation in richness when the minimum horizontal coverage was 20% for metagenomes from temperate latitudes (**Figure S14**).

Setting a horizontal genomic coverage threshold has an effect on the occurrence of each bin in the metagenomic samples. In all metagenomic datasets (Arctic, Southern Ocean and temperate), the distribution of occurrence vs mean abundance of bins stabilizes when the minimum horizontal coverage is 10% or higher (**Figure S15**). Lower thresholds show different patterns of distribution, increasing the number of higher occurrences at very low mean abundances (**Figure S15**). To date, there is no consensus about the minimum horizontal coverage thresholds to discard false mappings.

Based on our analyses, we chose 20% as the minimum horizontal genomic coverage to consider recruitments valid. Metagenomic read recruitments are accessible in Table S7 and metatranscriptomic read recruitments are in Table S8.

### Abundance and distribution of bins

#### Estimation of bin abundance and occurrence

Only those read recruitments aligning with an identity equal to or larger than 95% were considered to be representative of the bins. Recruitments passing the minimum horizontal genomic coverage threshold of 20% were considered to represent an actual presence of the bin in the sample. In comparison, those with a horizontal genomic coverage lower than 20% were considered not representative of the bin, thus absent in the sample. Read recruitments were transformed to RPKGs (recruited reads per genome kilobase and sample gigabase). Metagenomic RPKGs are accessible in Table S9 and metatranscriptomic RPKGs are in Table S10.

#### Distribution of communities based on MQ and HQ MAGs composition

The ordination of samples based on their MAG composition, with RPKG as an abundance estimate, was done with a Non-metric Multidimensional Scaling (NMDS) approach using function metaMDS from the vegan package in R.

### Niche breadth and classification of MAGs as specialists or generalists

#### Habitat specialist-generalist patterns in the Arctic Ocea

Specialist-generalist classification of MAGs was based on Levin’s Index (B)^67^. In order to avoid sampling biases, function spec.gen from R package EcolUtils (https://github.com/GuillemSalazar/EcolUtils) was used to calculate B for 1,000 random permutations of the metagenomic RPKG table and categorize MAGs into generalists if the original B index was larger than its confidence interval (CI 95) or specialists if the original B index was smaller than its confidence interval (CI 95). As the sampling occurred in a spatial and temporal gradient, each individual sample was considered as a habitat.

### Functional analysis of MAGs

To explore the ubiquity of representative biogeochemical cycling metabolisms related to carbon, sulfur, nitrogen and methane, a selection of 120 marker genes (**Table S2**) were searched in the Arctic MAGs dataset and only those pathways with enough encoded markers were considered valid.

### Phylogeny of RuBisCo large chain aminoacid sequences

A total of 14 RuBisCo large chain aminoacid sequences were detected by their KEGG Orthology annotation (K01601) in Arctic MAGs. They were aligned against the RuBisCo large-chain reference alignment profile published by ^52^ and the RuBisCo large-chain sequences from heterotrophic marine Thaumarchaeota published by ^93^ using Clustal Omega v1.2.3 (default options and 100 iterations) ^94^. Maximum-Likelihood phylogenetic reconstruction was done using the Jones-Taylor-Thorton model with FastTree v2.1.11 (default options) ^95^. Phylogenetic tree editing was done in iTol^96^.

### Definition of sentinel Arctic MAGs

Those MAGs that showed metagenomic recruitment exclusively in polar samples were selected and sentinel classification was done for the ones showing higher metatranscriptomic RPKGs per sample within each niche breadth category. For each sample and niche breadth category, all individual RPKGs were calculated relative to the highest in the group and only those RPKG recruitments representative of at least 50% of the highest RPKG recruitment per sample were selected as sentinels and shown in **Figure 7**.

## Acknowledgments

**General:** *Tara* Oceans (which includes both the *Tara* Oceans and *Tara* Oceans Polar Circle expeditions) would not exist without the leadership of the Tara Ocean Foundation and the continuous support of 23 institutes (http://oceans.taraexpeditions.org). We thank SHOOK Studio for assistance with figure design and execution.

## Funding

We further thank the commitment of the following sponsors and research funding agencies: the Spanish Ministry of Economy and Competitiveness (project MAGGY - CTM2017- 87736-R), CNRS (in particular Groupement de Recherche GDR3280 and the Research Federation for the study of Global Ocean Systems Ecology and Evolution, FR2022/Tara Oceans-GOSEE), European Molecular Biology Laboratory (EMBL), Genoscope/CEA, The French Ministry of Research, and the French Government ‘Investissements d’Avenir’ programmes OCEANOMICS (ANR-11-BTBR-0008), FRANCE GENOMIQUE (ANR-10-INBS-09-08), MEMO LIFE (ANR-10-LABX-54), PSL* Research University (ANR-11-IDEX-0001-02), ETH and the Helmut Horten Foundation, MEXT/JSPS/KAKENHI (projects 16H06429, 16K21723, 16H06437, 18H02279). SGA and CPA belong to the International Thematic Platform (PTI) Polar CSIC (https://polarcsic.es/en/). We also thank the support and commitment of Agnès b. and Etienne Bourgois, the Prince Albert II de Monaco Foundation, the Veolia Foundation, Region Bretagne, Lorient Agglomeration, Serge Ferrari, Worldcourier, and KAUST. The global sampling effort was enabled by countless scientists and crew who sampled aboard the Tara from 2009-2013, and we thank MERCATOR-CORIOLIS and ACRI-ST for providing daily satellite data during the expedition. We are also grateful to the countries who graciously granted sampling permissions. The authors declare that all data reported herein are fully and freely available from the date of publication, with no restrictions, and that all of the analyses, publications, and ownership of data are free from legal entanglement or restriction by the various nations whose waters the Tara Oceans expeditions sampled in. This article is contribution number XX of *Tara* Oceans.

## Notes

### Competing Interest Statement

The authors have declared no competing interest.

