## Supplementary Material for "Ecogenomics of key prokaryotes in the arctic ocean"

### Taxonomical classification of Arctic MAGs

Proteobacteria and Bacteroidota (previous Bacteroidetes) were the predominant bacterial phyla (182 and 111 MAGs, respectively, making up 34.3% and 20.9% of the detected diversity), which is in agreement with previous studies on microbial communities in the Arctic that were based on amplicon 16S iTAG sequencing (Pedrós-Alió et al., 2015). Thermoplasmatota (previously Euryarchaeota) was the dominant phylum of Archaea MAGs (50 MAGs; 9.4% of the total diversity). The most represented classes of these three dominant phyla were Alphaproteobacteria (61 MAGs; 11.5% of the total), Gammaproteobacteria (121; 22.8%), Bacteroidia (111; 20.9%) and Poseidoniia (50; 9.4%) (Figure 1D), all of which have been found previously to be present in Arctic waters. For example, the presence of Poseidoniia Archaea (or MGII) was reported from different regions of the Arctic (Bano et al., 2004), especially in the coast of Beaufort Sea, where waters are richer in labile organic matter of terrestrial origin (Galand et al., 2006). Alpha- and Gammaproteobacteria, as well as Bacteriodetes, are often the most abundant phyla in Arctic marine bacterioplankton (Pedrós-Alió et al., 2015), and their abundances seem to vary seasonally depending on sea ice dynamics (Pedrós-Alió et al., 2015). Other bacterial phyla less prevalent among our Arctic MAGs included Chloroflexota (previously Chloroflexi), Verrucomicrobiota (previously Verrucomicrobia), Actinobacterota (previously Actinobacteria), Planctomycetota (previously Planctomycetes) and Marinisomatota (previously Marinimicrobia or SAR406 clade). Multiple studies based on 16S rRNA genes have shown the presence of these phyla in polar waters (e.g., Bowman and McCuaig, 2003; Collins et al., 2010; Ghiglione et al., 2012; Storesund and Øvreås, 2013; Luria et al., 2016; Sipler et al., 2017). Archaeal MAGs included the Halobacterota (previously Euryarchaeota) and Crenarchaeota phyla (previously Thaumarchaeota). Crenarchaeota were found to dominate during the polar night (Alonso-Sáez et al., 2012) while Halobacterota, which dominated the archaeal assemblage of the deep Atlantic water masses from the central Arctic (Galand et al., 2009b), showed very low abundances throughout the year in surface waters of the Western Arctic (Kirchman et al., 2007; Alonso-Sáez et al., 2008).

SUPPLEMENTARY FIGURES

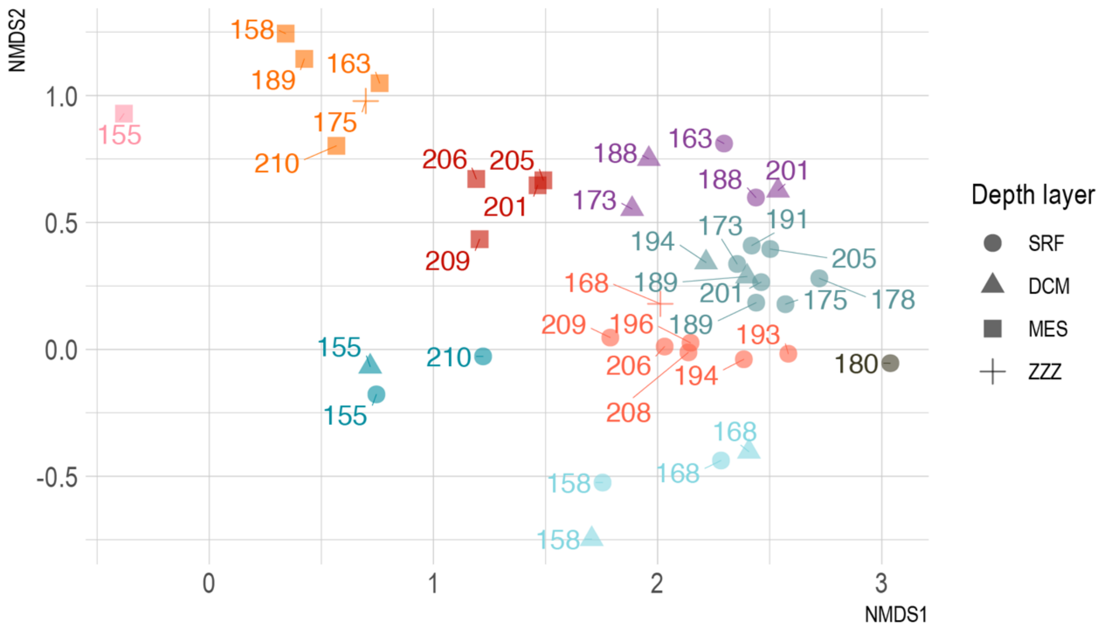


**Figure S1. NMDS of the 16S miTAG community composition of the 41 Tara Oceans Polar Circle metagenomes.**

Colors delimit the 9 groups of samples used for co-assembly in order to build Arctic bins. Shape indicates the ocean layer from which each metagenomic sample was collected.

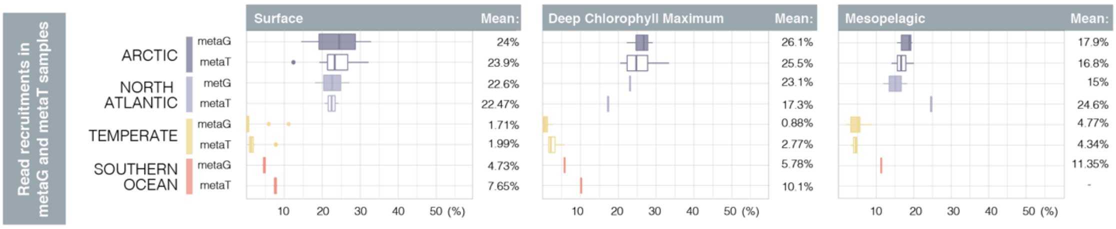


**Figure S2. Trends in metagenomic and metatranscriptomic read recruitments of a subset of 725 Arctic bins for comparison with Delmont et al. 2018 read recovery.**

Distribution of metagenomic (metaG) and metatranscriptomics (metaT) read recovery by the 725 bins that showed completeness >70% or assembly length >2Mbp, following Delmont et al., 2018 bin subsetting thresholds. Samples are divided by layer (columns) and latitudinal range (purple boxes for *Tara* Oceans Polar Circle, yellow boxes for temperate samples from *Tara* Oceans Expedition and red boxes for Southern Ocean samples from the *Tara* Oceans Expedition). Metagenomic samples are represented by filled boxplots, metatranscriptomic samples are represented by empty boxplots. Mean percentage of the read recruitments per group of samples is indicated at the right side of each plot.

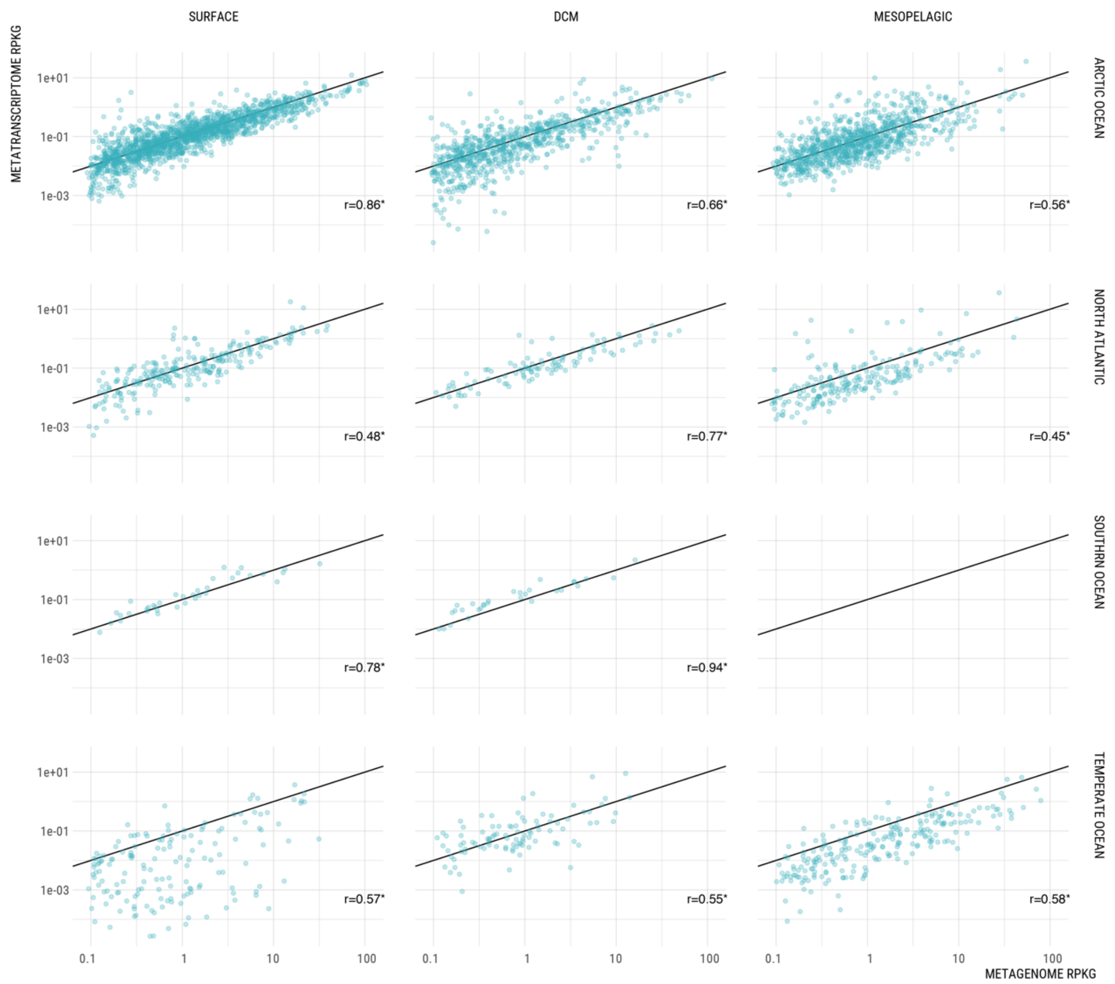


### Figure S3. Abundance vs activity recruitment plots in different latitudes.

Dotplots of metagenomic RPKGs (X axis) vs metatranscriptomic RPKGs (Y axis) of the 3550 individual bins, in the different layers (columns) and latitudes (rows). Significant Pearson's correlations (p-value <0.05) are shown with an asterisk.

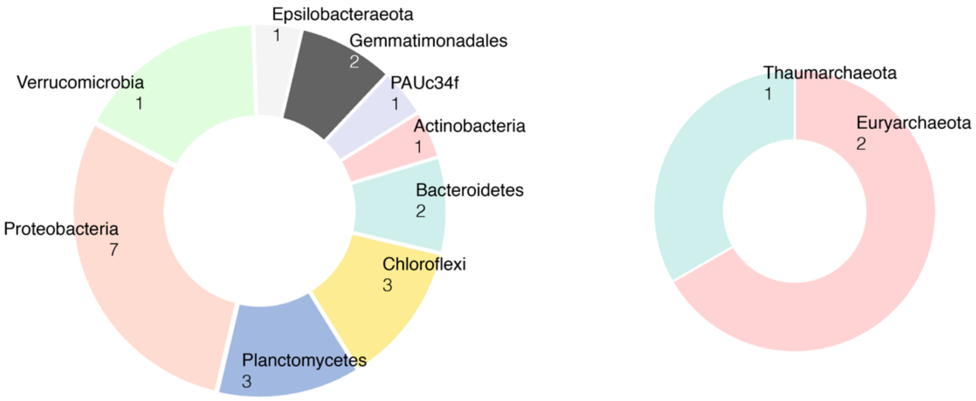


**Figure S4. Taxonomic classification of the 27 partial ribosomal genes encoded in the 530 MQ and HQ Arctic MAGs.**

A) Number of Arctic MAGs assigned to each phylum in the Bacteria domain
B) Number of Arctic MAGs assigned to each phylum in the Archaea domain

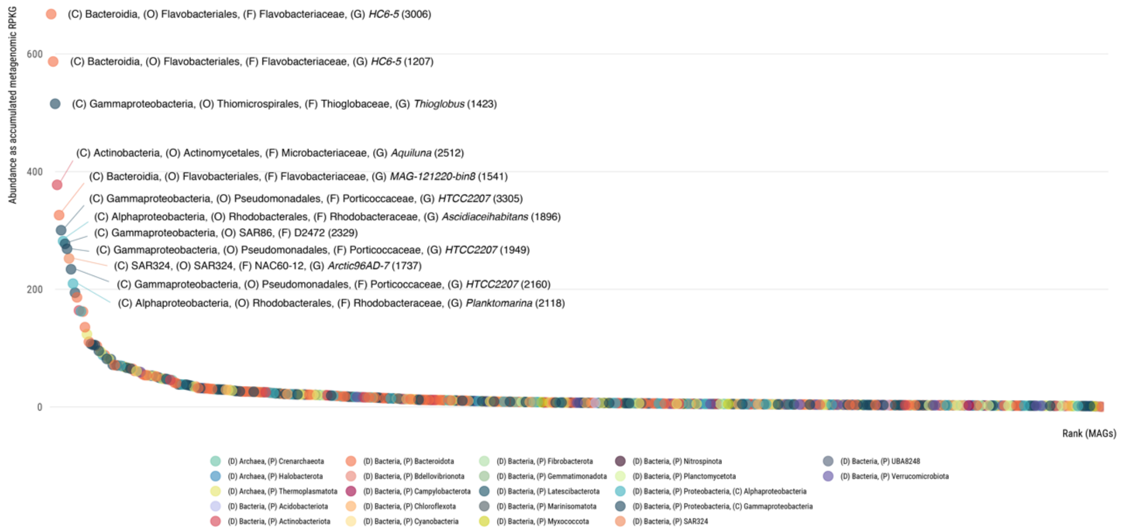


**Figure S5. Rank-abundance curve of Arctic MAGs in Arctic metagenomes.**

MAGs are sorted in X axis by their accumulated RPKGs in the 37 Arctic metagenomes (including the sub-Arctic North Atlantic) used in this study. MAGs are colored by phyla and the those recruiting at least 200 RPKGs are labelled with extended taxonomic annotation. Taxonomic annotation reaches the furthest level of classification for each MAG and the number in parenthesis is the MAG's identification code.

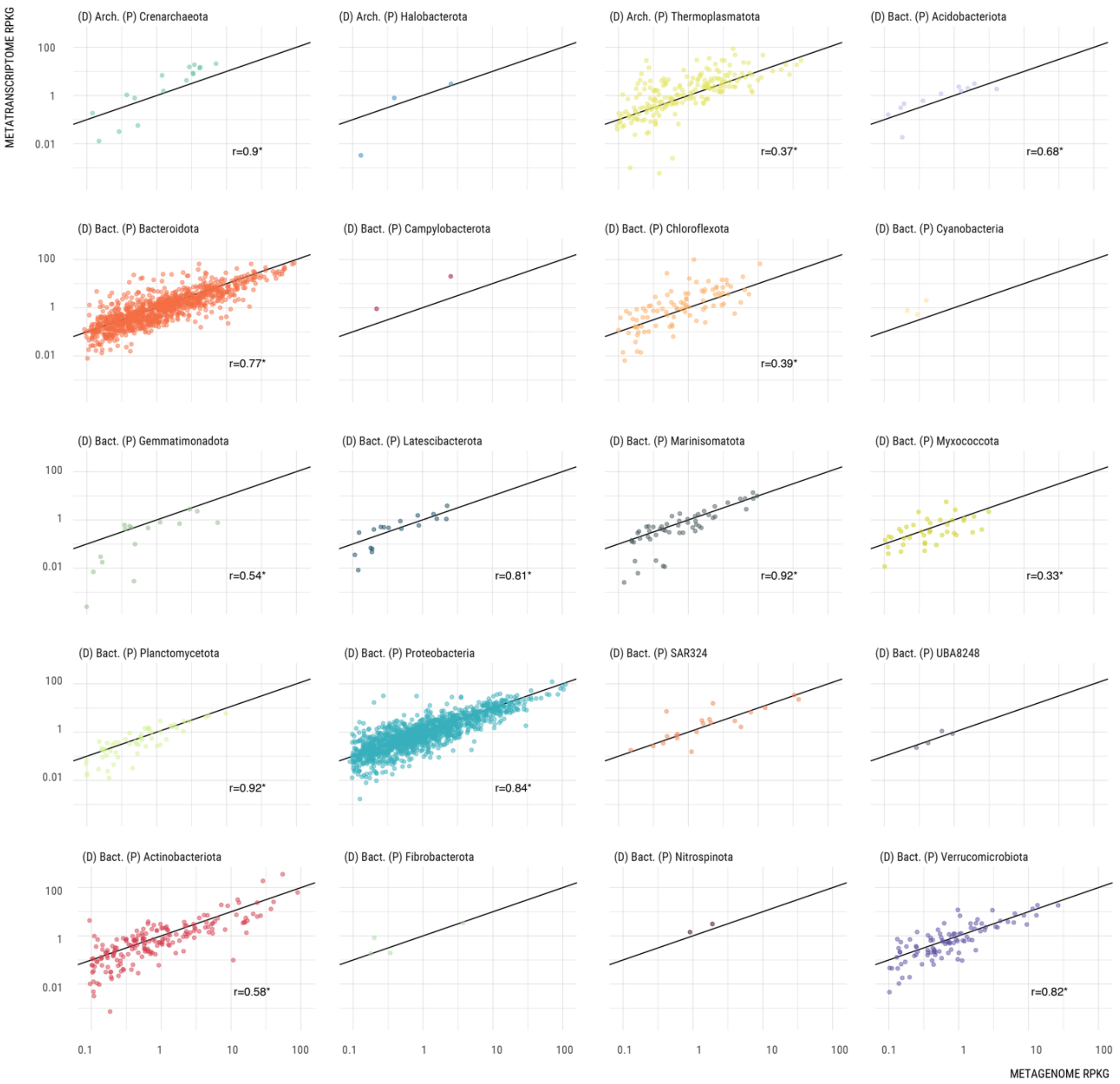


**Figure S6. Abundance vs activity recruitment plots of MAGs in the Arctic samples, by phyla.**

Comparison of the metagenomic RPKGs (X axis) vs metatranscriptomic RPKGs (Y axis) of the individual 530 Arctic MAGs separated by phylum, pooling all 37 Arctic samples together. Significant Pearson's correlations (p-value <0.05) are shown with an asterisk.

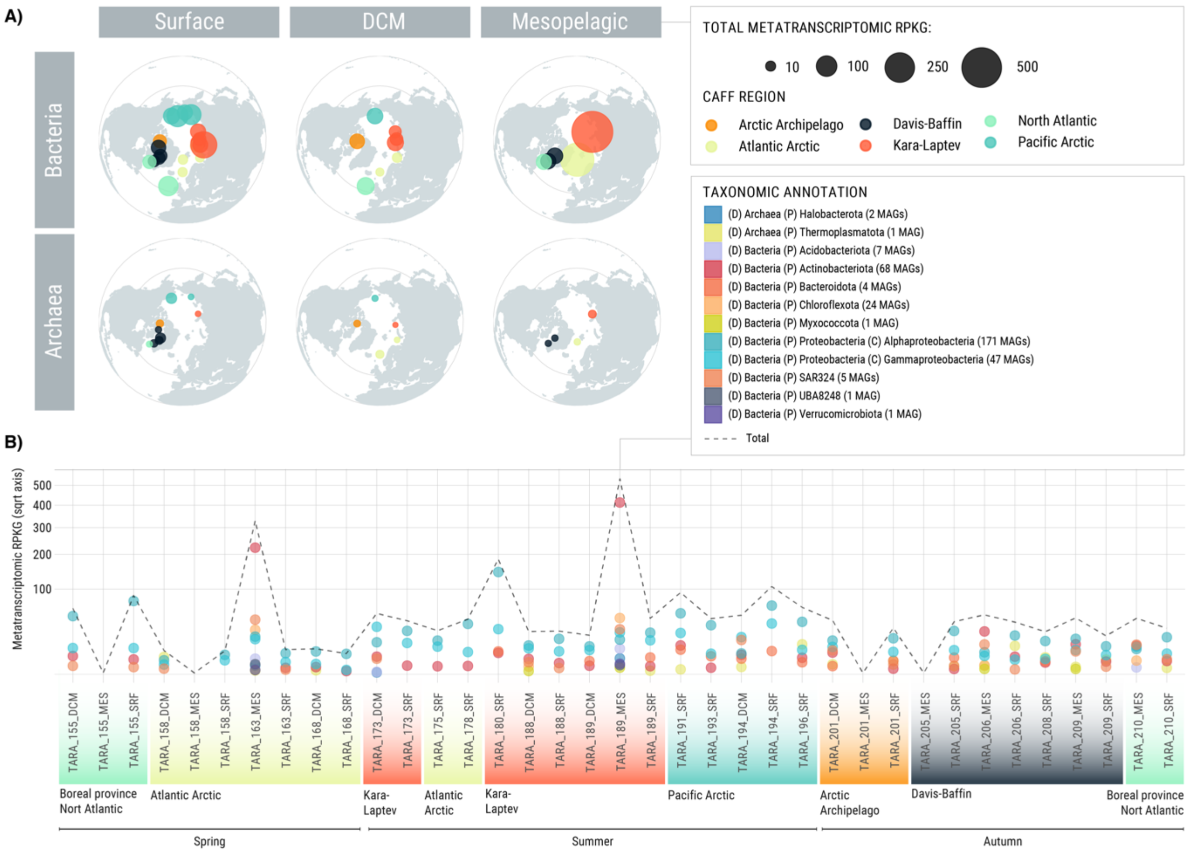


**Figure S7. Accumulated expression of the 332 Arctic MAGs containing the key marker gene *coxL* K03520 from the aerobic carbon-monoxide dehydrogenase.**

A) Polar maps with the accumulated metatranscriptomic RPKGs of 332 Arctic MAGs containing *coxL*, color-coded by CAFF region. The size of the dot is proportional to the accumulated metatranscriptomic RPKGs. Absent maps mean that no recruitment was found for that specific metabolism/domain/layer.
B) Metatranscriptomic RPKGs of 332 Arctic MAGs containing *coxL* colored based on taxonomic annotation at the phylum level. Accumulated RPKGs per sample is depicted with a dashed black line.

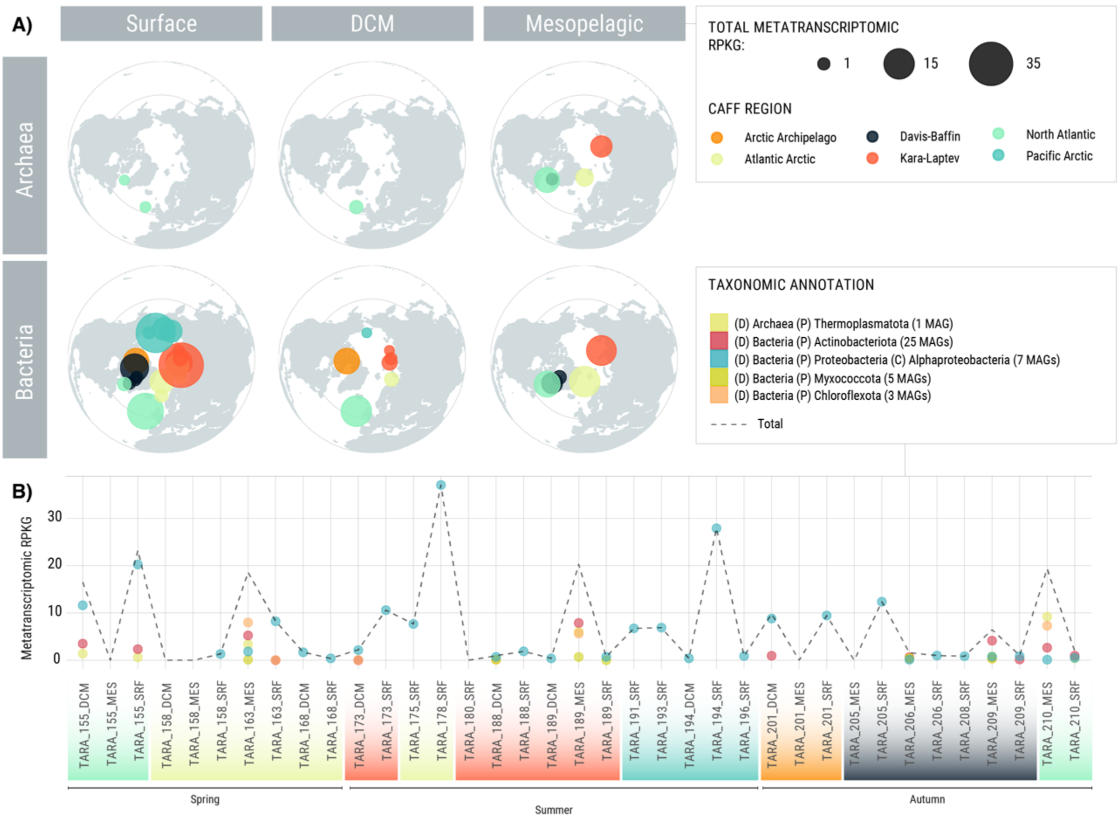


**Figure S8. Accumulated expression of 40 Arctic MAGs containing marker genes for the inorganic carbon fixation pathway 3-hydroxypropionate bicycle.**

A) all MAGs that contain at least one of the following key genes: for the 3-Hydroxypropionate bicycle: malyl-CoA/(S)-citramalyl-CoA lyase ( K08691) or propionyl-CoA carboxylase (K15052). The size of the dot is proportional to the accumulated metatranscriptomic RPKGs.
B) Metatranscriptomic RPKGs of 41 Arctic MAGs containing genes indicated above, colored based on taxonomic annotation at the phylum level. Accumulated RPKGs per sample is depicted with a dashed black line.

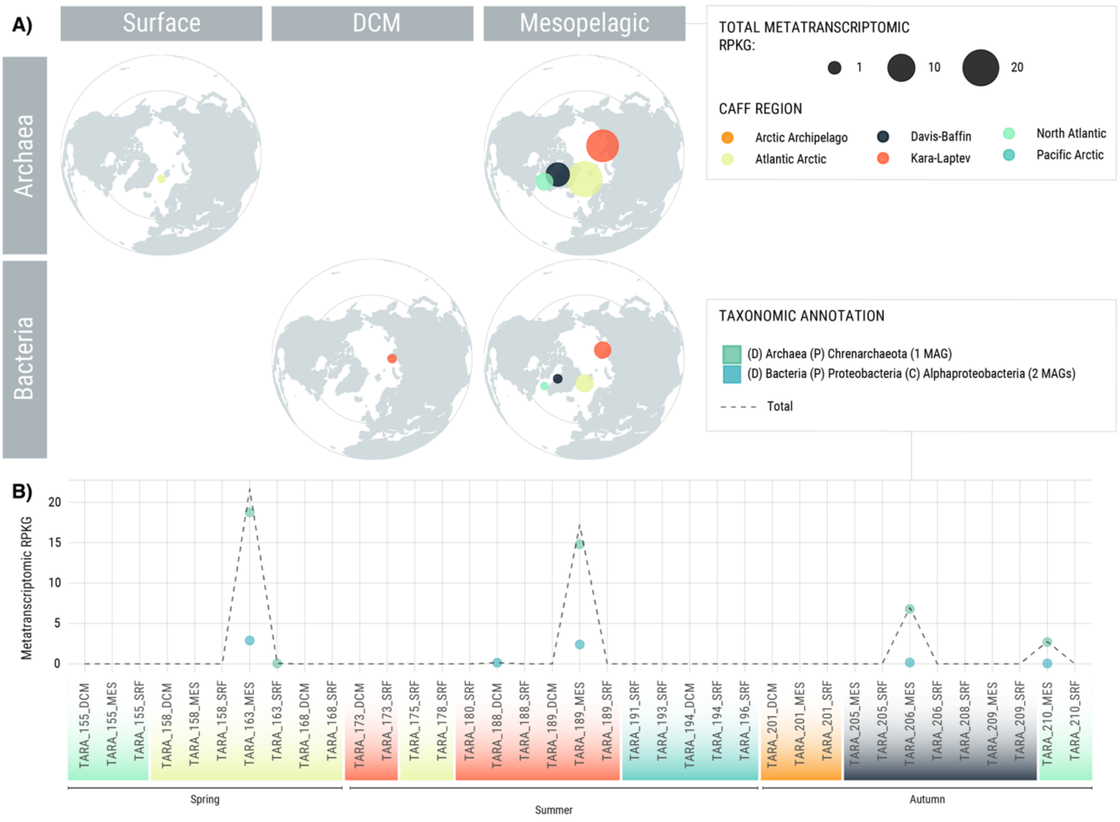


**Figure S9. Accumulated expression of 4 Arctic MAGs encoding marker genes for the inorganic carbon fixation pathway 3-hydroxypropionate/4-hydroxybutyrate cycle.**

A) Stacked bars with the accumulated metatranscriptomic RPKGs of all MAGs containing key enzyme 4-hydroxybutyryl-CoA dehydratase (K14534) together with the complementary gene 3-hydroxypropionyl-coenzyme A dehydratase (K15019). The size of the dot is proportional to the accumulated metatranscriptomic RPKGs.
B) Metatranscriptomic RPKGs of 3 Arctic MAGs containing genes indicated above, colored based on taxonomic annotation at the phylum level. Accumulated RPKGs per sample is depicted with a dashed black line.

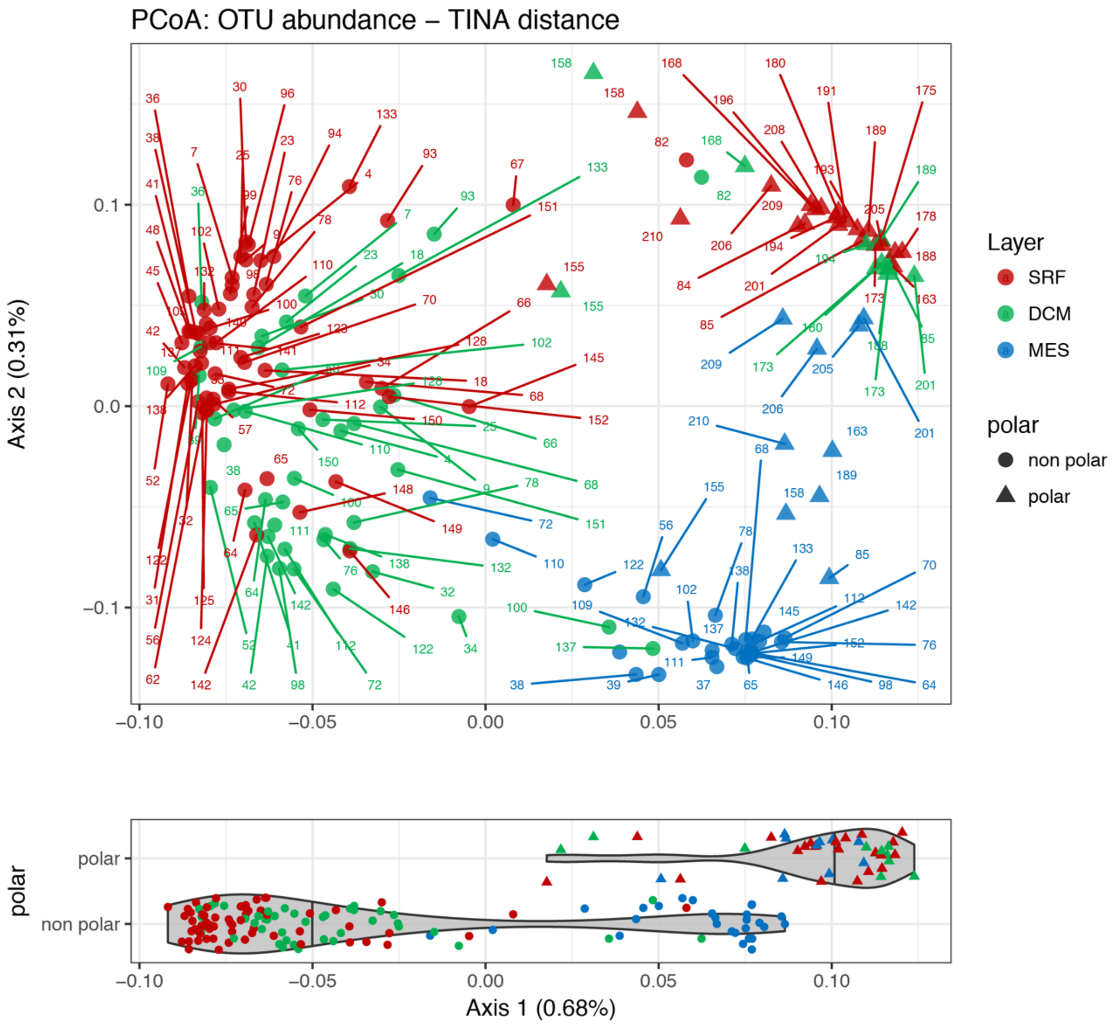


**Figure S10. PCoA of Tara Oceans expedition (2009-2013) samples including Arctic and sub-Arctic North Atlantic (polar) samples.**

PCoA based on TINA distances of miTAG community composition. Bottom violin plot splits distribution of polar and non-polar samples in axis 1.

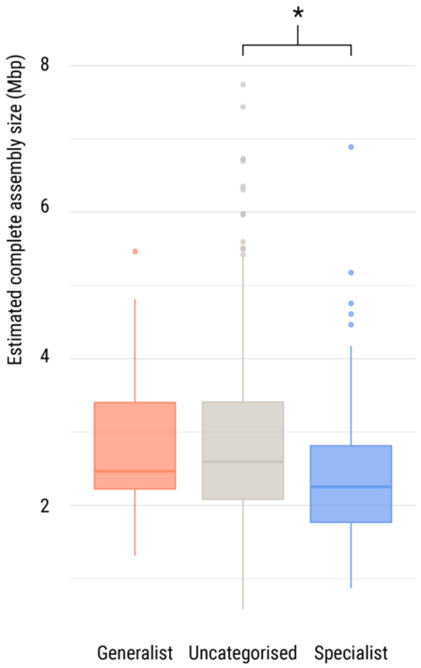


**Figure S11. Differences in estimated complete genome sizes.**

Between different niche breadth categories. DTK test shows significant differences (p-value < 0.05) between MAGs specific classified as habitat specialists and those uncategorized.

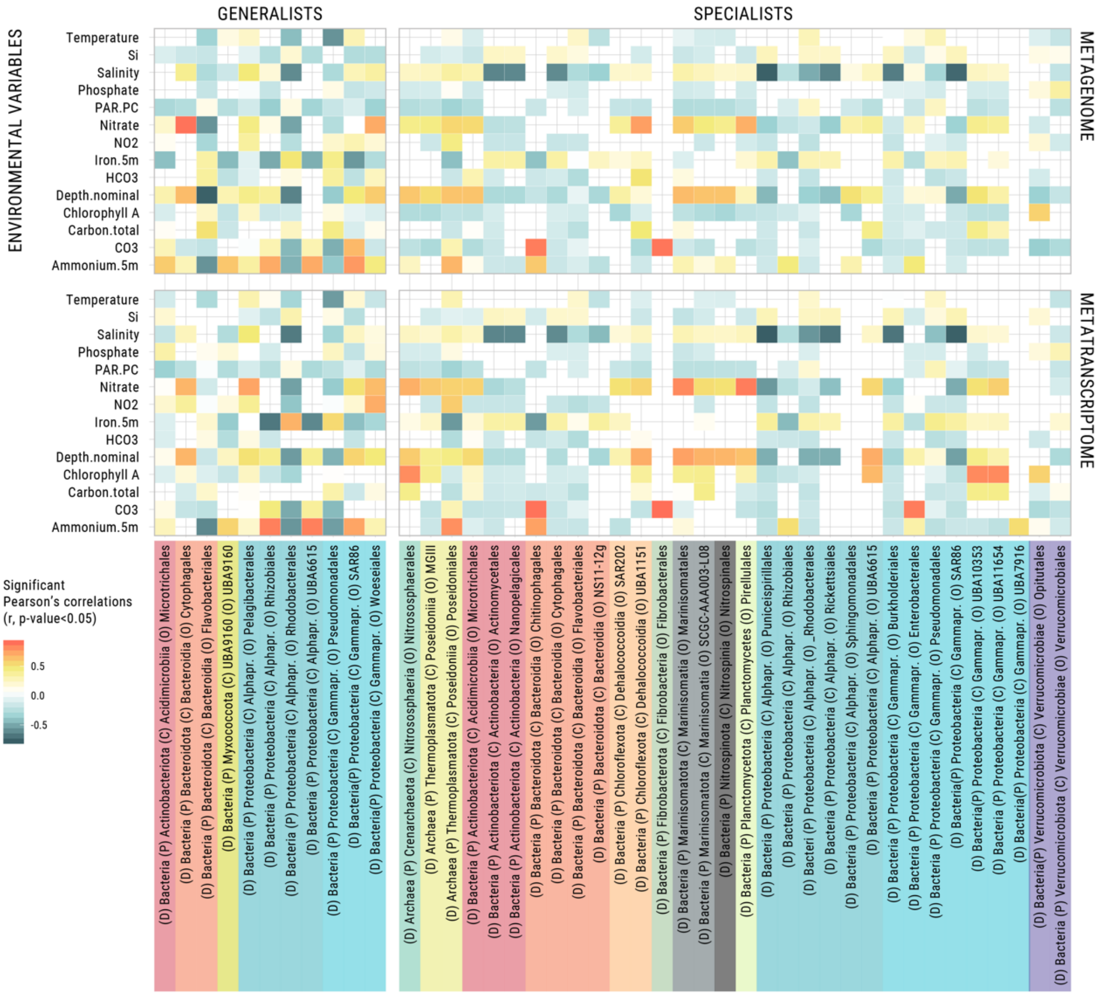


**Figure S12. Significant correlations between metagenomic and metatranscriptomic RPKGs of MAGs and environmental variables of Arctic sites do not show a key environmental parameter driving the abundance of either generalist or specialist MAGs.**

Significant Pearson's correlations (p-value <0.05) between accumulated metagenomic (top row) and metatranscriptomic (bottom row) RPKGs and environmental variables (Y axis). Metagenomic and metatranscriptomic RPKGs are accumulated within taxonomic orders. The left column contains correlations with generalist MAGs and right column contains correlations with specialist MAGs. Only significant correlations are shown.

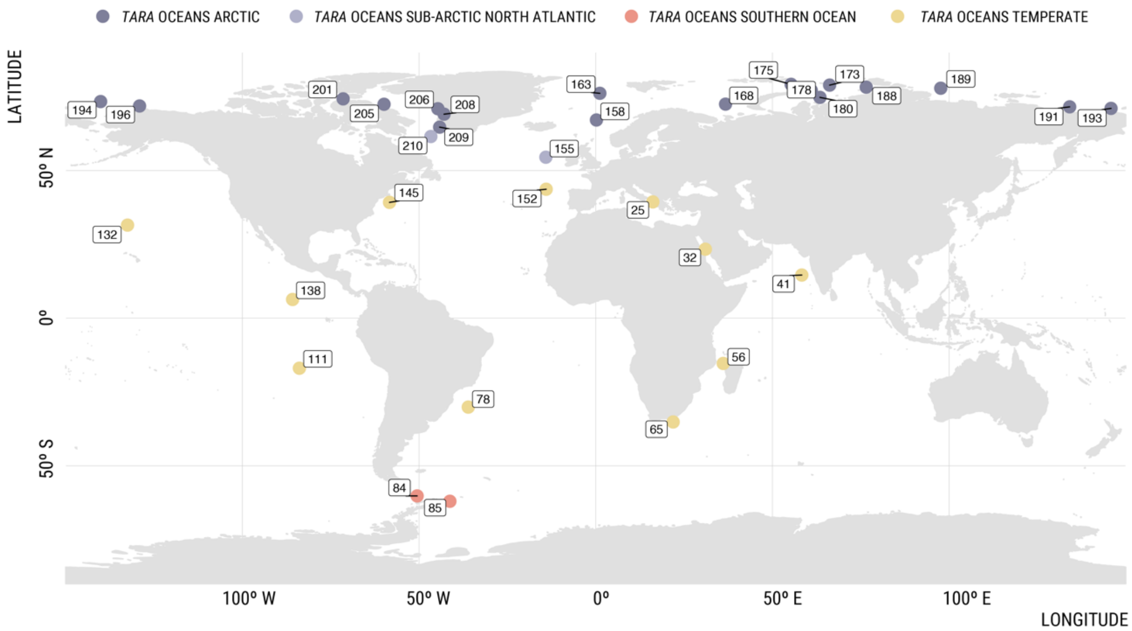


**Figure S13. Map with the reference stations with metagenomic and metatranscriptomic samples used in the study.**

Samples are colored based on the expedition. Table S4 contains more details about environmental metadata of these stations.

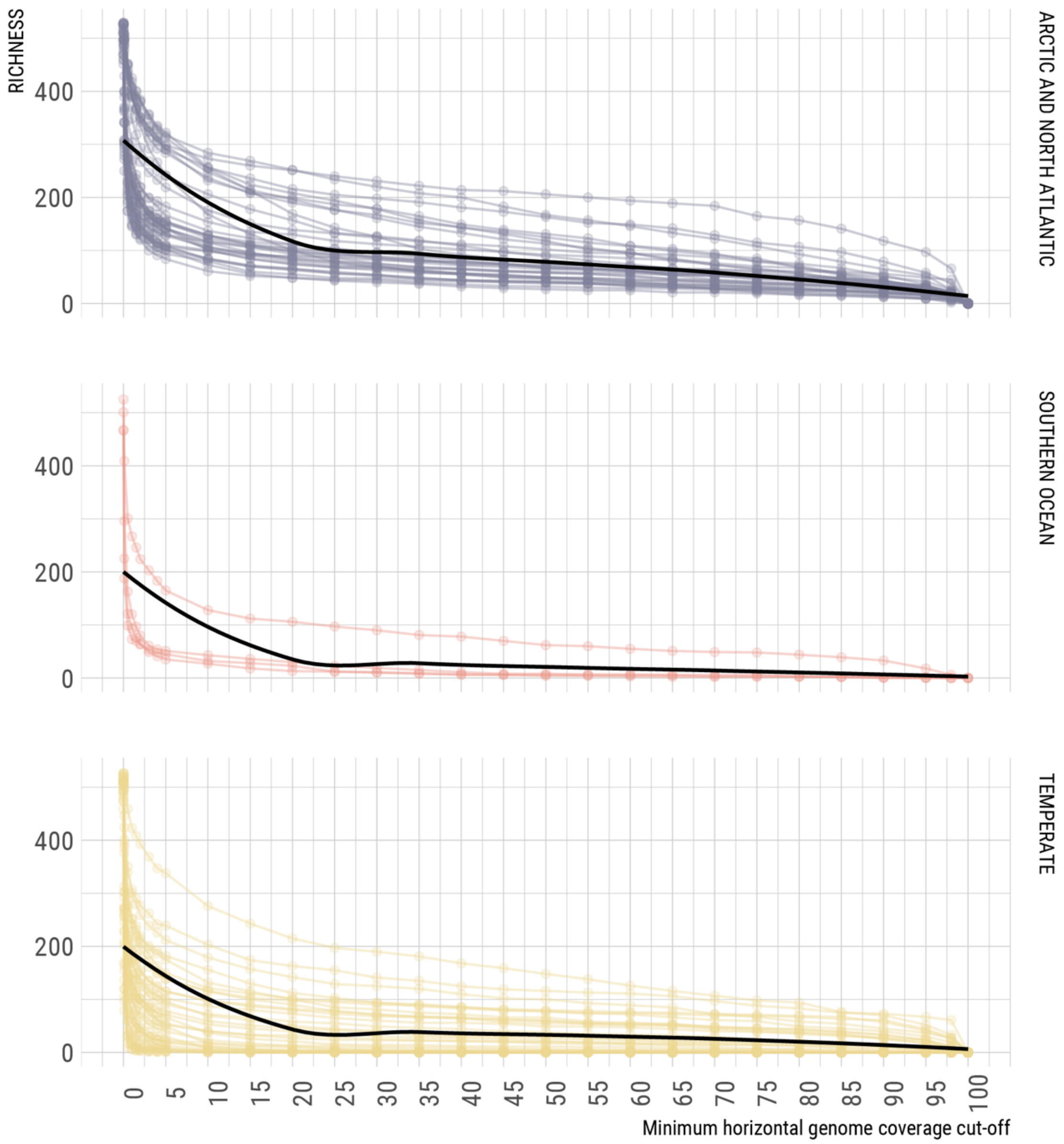


**Figure S14. Effect on samples' richness with increasing minimum horizontal coverage thresholds of metagenomic mappings.**

Purple data represents metagenomic samples from the Tara Oceans Polar Circle dataset (Arctic and sub-Arctic North Atlantic). Red data represents metagenomic samples from the Southern Ocean and yellow data represents metagenomic samples from temperate latitudes from the Tara Oceans Expedition.

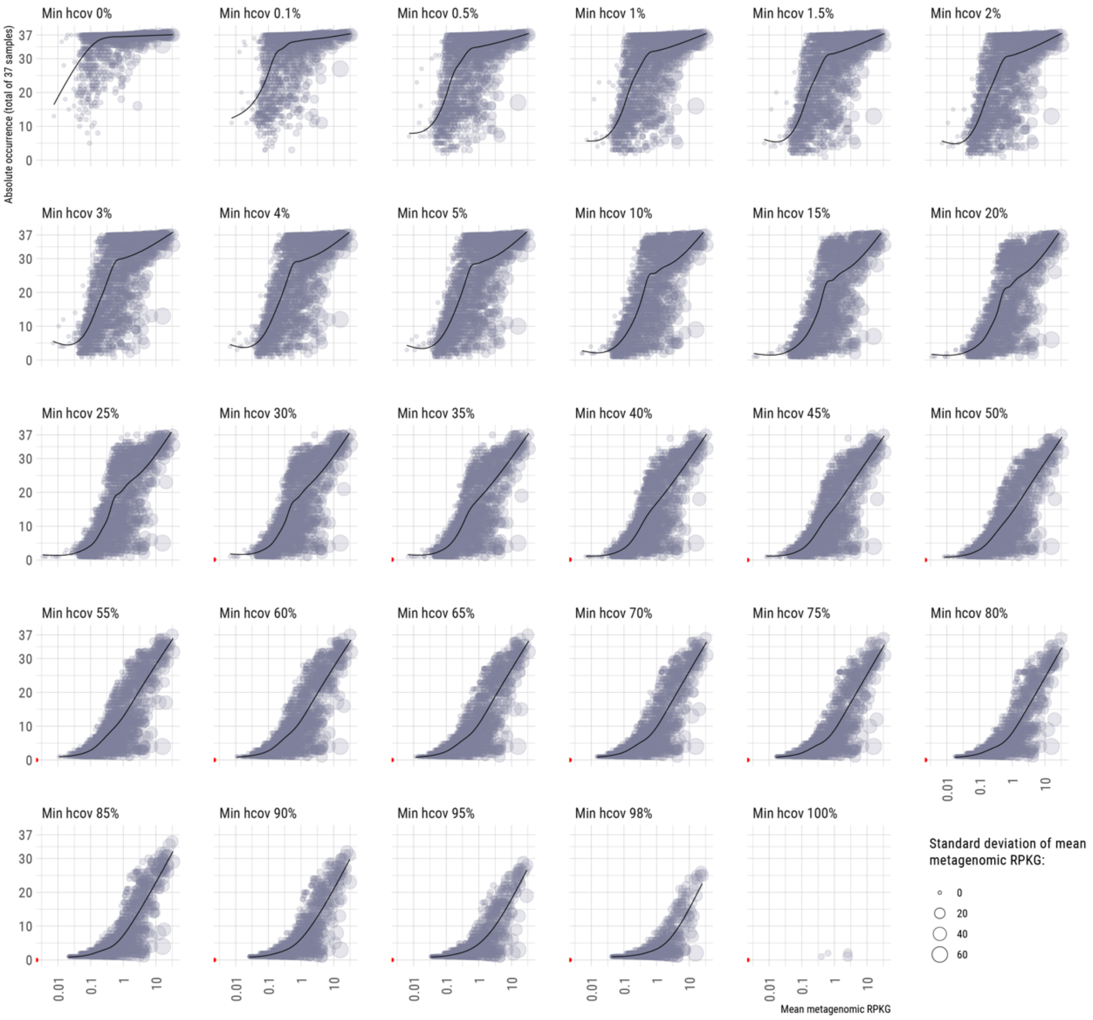


**Figure S15. Effect of increasingly stringent minimum horizontal coverages in MAG recruitments in *Tara* Oceans Arctic expedition samples.**

Each dot represents a MAG and its location in the X axis (mean metagenomic RPKG) and Y axis (occurrence in the 37 Arctic and Sub-Atlantic North Atlantic samples) varies with every minim horizontal coverage threshold applied to their recruitments (facets). Size of dot represents the standard deviation of the mean RPKG. Red dots indicate that mean abundance and occurrence equal 0.

SUPPLEMENTARY TABLES

Table S1 Bin information (taxonomy, metrics, biogeography, niche breadth)
Table S2 Marker KEGG KOs
Table S3 Mixotrophic RuBisCo coding MAGs
Table S4 Environmental metadata
Table S5 Pools of metagenomes for co-assembly
Table S6 Silva annotation of 16S rRNA gene sequences
Table S7 Metagenomic read recruitments
Table S8 Metatranscriptomic read recruitments
Table S9 Metagenomic RPKGs
Table S10 Metatranscriptomic RPKGs
